# Development of a Plant Growth Promoting Bacterial EcoBiome Derived from Desert Soil Isolates

**DOI:** 10.1101/2025.07.02.662819

**Authors:** Camila Albarrán-Cuitiño, Daniel E. Palma, Hugo González, Mauricio González, Alexis Gaete

## Abstract

The application of plant growth-promoting (PGP) bacteria is increasingly studied for its potential to improve plant tolerance to biotic and abiotic stress. Developing synthetic microbial consortia represents a promising strategy, as it can enhance colonization success and functional synergy within the rhizosphere. In this study, we designed a stable EcoBiome derived from a synthetic community (SynCom) of 17 bacterial isolates obtained from three desert environments. We evaluated their PGP traits, including siderophore production, indoleacetic acid (IAA) synthesis, phosphate solubilization, and nitrogen fixation. Using Oxford Nanopore Technologies (ONT) sequencing of 16S rRNA genes, we tracked changes in relative abundance across successive subcultures under four temperature conditions. From this analysis, *Erwinia rhapontici* 1SR, *Pseudomonas yamanorum* RZ5, and *Plantibacter* sp. RU18 were identified as the dominant isolates and subsequently selected to construct the EcoBiome. Functional characterization showed that these isolates exhibited complementary PGP traits, biofilm formation capacity, and tolerance to water stress, both individually and in combinations. These findings highlight the potential of desert-derived bacterial consortia as microbial resources for developing biostimulants to enhance plant resilience under environmental stress conditions.

**Importance:** Several studies have focused on obtaining bacterial isolates with plant growth promoting traits, however, their success as microbial inoculants with plant stimulant activity is diminished (or null) because they do not have the ability to compete efficiently with the natural soil microbiome. Thus, in this study we designed a synthetic community of 17 bacteria from desert environments and individually characterized their plant growth-promoting attributes, and in parallel we subjected this synthetic community to co-culture, subsequently evaluating its temporal prevalence (24 and 48 hours of culture) and at four different temperatures. With this, we developed and presented a stable EcoBiome of three isolates (one from each desert), stable over time, with attributes that promote plant growth and proliferation in hostile conditions, such as drought, with the purpose of being used as microbial inoculants as a whole, capable of competing, proliferating and forming part of the rhizosphere microbiome.

## 1. Introduction

The plant-associated soil bacterial community has garnered increasing attention in recent years due to its potential to enhance plant resilience against a variety of biotic and abiotic stresses (Bardgett and Caruso, 2020; Kumar and Verma, 2018; Wang et al., 2023). As sessile organisms, plants depend primarily on their root systems for essential biological processes and interactions. Symbiotic association between plants and bacteria within the rhizosphere have been shown to improve adaptive capacity by modulating key physiological processes, including nutrient acquisition, pathogen control, growth promotion, and increased tolerance to environmental stressors (Devi et al., 2022; Pascale et al., 2020; Tian et al., 2023). These effects, whether direct or indirect, significantly influence plant development. Such bacteria are classified as Plant Growth Promoting (PGP) organisms, with their role in plant health and productivity remaining an active field of research.

Given the beneficial properties of PGP bacteria, they have been proposed as promising alternatives to synthetic fertilizers and agrochemicals (Backer et al., 2018; Yadav et al., 2023), particularly through exogenous inoculation with isolated strains or polymicrobial consortia. This strategy has been tested in diverse crop species, including tomato (*Solanum Lycopersicum)* (Katsenios et al., 2021; Subramanian et al., 2016; Yavarian et al., 2021; Zuluaga et al., 2021), wheat (*Triticum aestivum*) (Akhtyamova et al., 2023; Khan et al., 2022; Mirskaya et al., 2022), and several fruit-bearing plants (Bokszczanin et al., 2021; Diniz et al., 2025; Ge et al., 2022; Gopalakrishnan et al., 2022; Karlidag et al., 2007; Solórzano-Acosta and Quispe, 2024). These studies emphasize the critical role of microbial diversity in maintaining and enhancing ecosystem functions.

Resent research suggests that polymicrobial consortia offer superior benefits over single-strain inoculants, primarily due to synergistic functional interactions that can more effectively promote plant growth. For example, inoculation with multiple PGP strains has been associated with improved plant biomass accumulation (Wang 2022) and increased shoot and root growth in walnut seedlings (Kaur 2021). However, designing effective synthetic microbial communities remains challenging. Antagonistic interactions between bacterial strains, competitive exclusion, and limited tolerance to environmental stress can compromise their stability and functionality (Mehrabi et al., 2016; Wang et al., 2022).

In this context, microbial communities native to desert soils exhibit a remarkable metabolic versatility and adaptive capacities compared to those from controlled environments or less stresful environments (Fierer et al., 2012; Mandakovic et al., 2023; Neilson et al., 2012; Palma et al., 2025). These microorganisms can thrive under extreme conditions, including water scarcity, high salinity, and UV radiation (Gao and Garcia-Pichel, 2011; Marasco et al., 2022). Therefore, exploring desert-derived PGP bacteria represents a promising strategy to mitigate environmental stress in agriculture (Alsharif et al., 2020; Gaete et al., 2021). Furthermore, these bacteria often display a strong capacity colonize root tissues and may even be vertically transmitted through seeds (Jooste et al., 2019).

Several studies have focused on isolating soil bacteria from desert environments with PGP attributes (Cherif et al., 2015; Gaete et al., 2020; Yadav et al., 2015), highlighting their potential as inoculants to enhance plant resilience under biotic and abiotic stresses (Alsharif et al., 2020; Chen and Costanza, 2024; Kochhar et al., 2022). However, challenges remain in designing stable and functional synthetic communities from these native microorganisms.

In this study, we present a workflow for constructing a stable EcoBiome, defined here as a synthetic microbial community composed of persistent bacterial members following successive subcultures. We isolated and characterized 17 bacteria strains from desert soils in Chile. These bacteria were analyzed through molecular and culture-based techniques to confirm their PGP potential, including tolerance to water deficit conditions. This study provides a functionally validated desert-derived EcoBiome with PGP attributes and tolerance to stress, contributing to the development of microbial resources for sustainable agriculture.

## 2. Materials and Methods

### 2.1. Sampling Sites and Isolation of Bacterial Strains

A total of 17 bacterial isolates were obtained from three sites classified as desert environments (Figure 1), based on the criterion of annual precipitation below 250 mm/year (Cheng et al., 2017). Soil samples were collected from Atacama Desert (23°21’06.9"S 67°50’01.0"W) in 2022, from the Experimental Station “Las Cardas” in the Limarí Valley, Coquimbo (30°15’06.6"S 71°15’23.8"W) in 2019, and from a site near the Union Glacier Station, Antarctic Desert (79°46’51.6"S 83°18’50.1"W) in 2022. At each location, surface soil was removed, and soil samples were collected from a depth of up to 20 cm. Samples were stored in sterile bags and kept refrigerated at 4°C until further processing.

**Figure 1.**
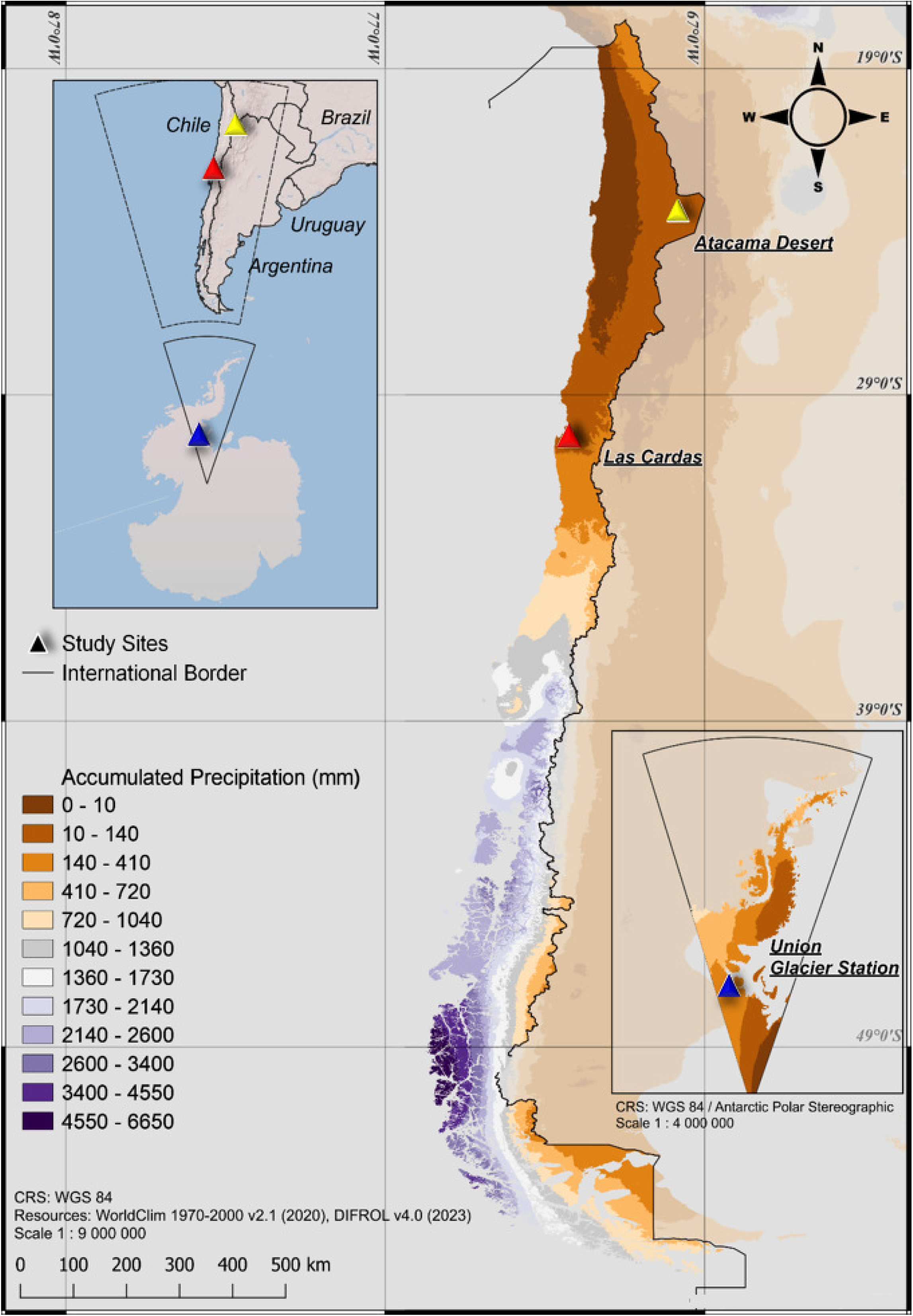
Map of annual precipitation in Chile. Colored triangles indicate the sampling sites included in this study: the Atacama Desert (Yellow), the “Las Cardas” Experimental Station in the Limarí Valley (Red) and the Antarctic peninsula (Blue).

### 2.2. Bacterial Isolation and Taxonomic Identification via 16S rRNA Gene Sequencing

Isolation was performed as per Mandakovic et al. (2018). Briefly, 1 g of each soil sample was thoroughly mixed with 1 mL of phosphate buffer saline (PBS) for 2 hours using a Revolver Rotator (Labnet). After centrifugation at 8,000 rpm for five minutes, 100 uL of supernatant was plated onto a Luria Bertani (LB) agar and incubated at 10°C, 20°C, and 30°C for 72 hours. Individual colonies were cultured in 24-well plates with 1 mL of LB medium, shaken at 160 rpm at 25°C for 24 hours. The resulting cultures were stored at -80°C with glycerol in the internal microbial repository.

Genomic DNA was extracted using the DNeasy Blood & Tissue Kit (Qiagen) following the manufacturer’s specifications. The 16S rRNA gene was amplified with universal primers 27F/1492R at 10µM, 2µL of genomic DNA, 12.5µL of 2X GoTaq® G2 Green Master Mix (Promega), and 8.5 µL of nuclease-free water, in a final volume of 25 µL. PCR conditions consisted of an initial denaturation at 95°C for 3 minutes, followed by 30 cycles of denaturation 95°C for 30 seconds, annealing at 61°C for 30 seconds, and extension at 72°C for 90 seconds. A final extension step was performed at 72°C for 10 minutes.

Amplicon was sequenced using Sanger sequencing (Macrogen) and trimmed using the CLC Genomics Workbench (Qiagen). Taxonomic identification was conducted using the EzBioCloud 16S database, and all sequences were deposited in the National Center for Biotechnology Information (NCBI) Genbank database (Table S1).

### 2.3. In Vitro Characterization of Plant Growth-Promoting Traits

Plant growth-promoting (PGP) assays were performed as described in Gaete et al. (2020), with slight modifications. Briefly, different culture media were used to assess four PGP traits. Bacterial isolates were incubated at 30°C for five days under static conditions.

Phosphate solubilization and nitrification were evaluated using media prepared according to the technical specifications provided by HIMEDIA. Pikovskaya agar (PKV) was used to assess phosphate solubilization (Dhull et al., 2018), while a modified nitrification medium (NM) agar was used to test for nitrification (He et al., 2016). In both cases, the formation of a clear halo around bacterial colonies was interpreted as a positive result.

Siderophores production was assessed using chrome azurol S (CAS) agar medium, following the method described by Louden et al. (2011). A color change from blue to orange, accompanied by the formation of a halo around the colony, was interpreted as a positive result. The diameter of the halo was recorded as a proportional measure of siderophore production.

The production of indole-3-acetic acid (IAA) was determined by a colorimetric assay using Salkowski reagent, following the method described by Mehmood et. al. (2021). A color change from yellow to red indicated IAA synthesis. The intensity of the color change was quantified by measuring absorbance and compared against a standard curve prepared with known IAA concentrations (0, 5, 10, 20, 50 and 100 µg/mL).

All statistical analyses were performed using GraphPad Prism software version 8.0.1 for Windows (GraphPad Software, La Jolla, CA, USA; www.graphpad.com).

### 2.4. Design and Evaluation of a Synthetic Co-Culture

Each bacterial isolate was independently cultured in LB medium at 15°C, 20°C, 25°C, and 30°C for 24 hours with shaking at 160 rpm (Figure 2). The optical density (OD_600_) of each culture was measured and adjusted to 0.1 using an Infinite 200 PRO NanoQuant spectrophotometer (Tecan), in a final volume of 10 mL. The culture was then divided into two equal fractions of 5 mL each. One fraction was used for genomic DNA extraction at time zero (t₀), and the second was used to evaluate the influence of temperature within an agronomic range (15 °C to 30 °C) on bacterial growth in culture. Two subsequent subcultures (t₁ and t₂) were performed at 24-hour intervals, using a 1:10 dilution (500 µL of bacterial culture into 5 mL of fresh LB medium) (Figure 2). After each incubation period, DNA was extracted from the synthetic community using the E.Z.N.A.® Soil DNA Kit (Omega Bio-Tek), following the manufacturer’s instructions.

**Figure 2.**
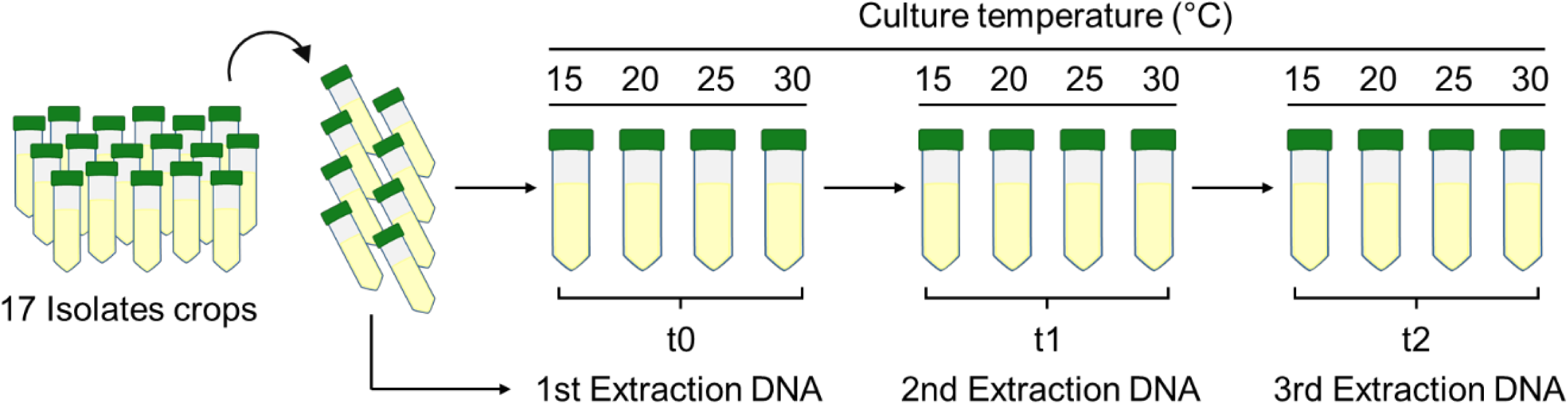
Schematic representation of the bacterial co-culture method at different temperatures and DNA isolations time points. The first DNA extraction (t₀) was performed at the start of the co-culture of the 17 isolates, followed by two subcultures after 24 hours of incubation each (t₁ and t₂).

### 2.5. Amplicon Sequencing and Diversity Analysis of the Synthetic Community

The extracted DNA was used to analyze the bacterial diversity through amplification of the 16S rRNA gene using the 16S Barcoding Kit SQK-RAB204 (Oxford Nanopore Technologies), following the manufacturer’s instructions. Sequencing was performed on a MinIon device version 1B (Oxford Nanopore Technologies) using a R9.4.1 flow cell.

Basecalling and demultiplexing of reads were conducted with Dorado v0.5.2 (Oxford Nanopore Technologies), using the model dna_r9.4.1_e8_sup@v3.6. Filtering criteria included a minimum mean Q-score of 7, a minimum read length of 1,000 bp, and maximum read length of 2,000 bp. Read quality control was assessed using pycoQC v2.5.2 (Leger and Leonardi, 2019). Consensus 16S rRNA sequences for the bacterial isolates were generated using NanoCLUST (Rodríguez-Pérez et al., 2021).

For each isolate, a single 16S rRNA sequence was selected according to the following priority order: (1) genomic-derived sequence, (2) NanoCLUST consensus sequence, or (3) Sanger-derived sequence. Selected sequences were aligned using Infernal v1.1.4 (Nawrocki and Eddy, 2013) with the RF00177 covariance model from Rfam (Kalvari et al., 2020) and manually trimmed using AliView v1.28 (Larsson, 2014) to ensure comparable sequence lengths across isolates. Finally, Emu v3.4.5 (Curry et al., 2022) was used to estimate the relative abundances of each isolate in the synthetic community using a custom database constructed from the selected 16S rRNA sequences.

The phylogenetic Neighbor Joining tree was constructed using the 16S rRNA sequences of each bacterial isolated. The sequences were trimmed using the CLC software (QIAGEN Bioinformatics), multiple aligned and edited in the BioEdit biological sequence alignment editor (Hall, T.A., 1999) and finally clustered and visualized as a phylogenetic tree based on the neighbor joining technique using 1,000 replicates as bootstrap to obtain statistical support performed on MEGA software (Tamura, K et al., 2021).

### 2.6. Genomic DNA Extraction, Whole-Genome Sequencing, Assembly, and Annotation

Whole genome sequencing was performed using the previous extracted DNA in Section 2.5. Library preparation was conducted using the Ligation Sequencing Kit (Oxford Nanopore Technologies), following the manufacturer’s protocol, and sequencing was performed on a MinIon device (version 1B) with an R9.4.1 flow cell. The bioproject is available at the National Center for Biotechnology Information (NCBI) under the number PRJNA999330 and each genome identifier (ID): *Erwinia rhapontici* 1SR (GCA_050613375.1), *Plantibacter* sp. RU18 (GCA_050613355.1) and *Pseudomonas yamanorum* RZ5 (GCA_050613365.1).

Basecalling of Nanopore reads was performed with Dorado v0.5.2 (Oxford Nanopore Technologies) using the model dna_r9.4.1_e8_sup@v3.6, applying the following criteria: a minimum mean Q-score of 7 and a minimum read length of 1,000 bp. Read quality was assessed using pycoQC v2.5.2 (Leger and Leonardi, 2019). Consensus genome assemblies were generated with Trycycler v0.5.4 (Wick et al., 2021), based on 12 individual assemblies from read subsets: four generated using Raven v1.8.1 (Vaser and Sikic, 2021), four with Flye v2.9.2 (Kolmogorov et al., 2019), and four using the combination of Minimap2 v2.26 (Li, 2018), Miniasm v0.3 (Li, 2016), and Minipolish (Wick and Holt, 2019). The resulting consensus contigs were polished using Medaka (Oxford Nanopore Technologies). Circular contig were rotated to start at the dnaA gene, when present, using the fixstart function of Circlator v1.5.5 (Hunt et al., 2015). Genome completeness and contamination were assessed using CheckM v1.2.2 (Parks et al., 2015) with the lineage workflow. Taxonomic classification was performed using GTDB-Tk v2.3.2 (Chaumeil et al., 2020), employing GTDB release 214 (Parks et al., 2022). Genome annotation was conducted with Bakta v1.8.1 (Schwengers et al., 2021), using database version 5.0. Circular genome plots were generated using Circos (Krzywinski et al., 2009). Genome sequence were uploaded to the Type (Strain) Genome Server (TYGS) for a whole- genome-based taxonomic analysis (Meier-Kolthoff and Göker, 2019), incorporating recent methodological improvements (Meier-Kolthoff et al., 2022). Nomenclature information, synonymy and relevant taxonomic literature were retrieved via TYGS’s sister database, the List of Prokaryotic names with Standing in Nomenclature (LPSN) (Meier-Kolthoff et al., 2022). A phylogenetic tree was constructed using FastME 2.1.6.1 (Lefort et al., 2015), based on genome BLAST distance phylogeny (GBDP) distances calculated from genome sequences. Branch lengths were scaled according to GBDP distance formula d5. Bootstrap support values (>60%) from 100 replications are indicated above the branches, with an average support of 94.9%. The tree was rooted at the midpoint (Farris, 1972).

### 2.7. Genome-Based Prediction of Metabolic Pathways and Secondary Metabolites

A comprehensive analysis of metabolic pathways and secondary metabolite biosynthesis was conducted using KofamKOALA and antiSMASH, respectively. KofamKOALA tool utilizes a reliability-based K-value scoring system (Aramaki et al., 2020) to compare protein sequences with KEGG (Kyoto Encyclopedia of Genes and Genomes) orthologs, allowing the identification of metabolic pathways present in the bacterial genomes. Particular emphasis was placed on detecting KEGG orthologs associated with siderophore production, tolerance to heat and water stress, IAA biosynthesis, nitrogen fixation, 1-aminocyclopropane-1-carboxylate deaminase (ACCd) activity, and iron and phosphate uptake, as described by Gaete et al. (2020) and Chávez-Ávila et al. (2023). The analysis was also extended to identify genes associated with biofilm formation, as well as the biosynthesis of pyocyanin and pyoverdine.

The biosynthetic potential for specialized metabolites was assessed using antiSMASH version 7.0 (Blin et al., 2023) for bacterial genomes. This tool identifies biosynthetic gene clusters (BGCs), and analyses were conducted using the "relaxed" stringency parameter to maximize the detection of potential clusters.

### 2.8. EcoBiome Assembly and Compatibility Assessment

The EcoBiome was constructed following the definition and selection criteria described by Mousa et al. (2024). Briefly, the selection process aimed to preserve the representativeness of native bacterial communities from each desert environment, as well as their plant growth-promoting (PGP) traits. Additionally, an abundance criterion under co-culture conditions was applied to select isolates that demonstrated greater persistence after successive subculturing. Based on these criteria, three bacterial isolates were selected: *Erwinia rhapontici* 1SR, *Pseudomonas yamanorum* RZ5, and *Plantibacter* sp. RU18, which were used to evaluate the establishment of the EcoBiome.

Compatibility between the selected isolates was assessed through competition and attraction assays. The competition activity was performed on LB agar medium following the procedure described by Pérez-y-Terrón et al. (2014), while the attraction activity was carried out according to Berendsen et al. (2018). For the competition assay, 100 µL of each isolate was inoculated onto the surface of a Petri dish and allowed to dry for five minutes. Subsequently, 10 uL of a second isolate was inoculated at the same location. Inhibition was indicated by the formation of a clear halo around the bacterial culture. The attraction assay involved inoculating 1 µL of each individual isolate diagonally in V-shape rows. The assay was considered positive when the bacterial colonies exhibited chemotactic behavior, evidenced by changes in their circular morphology as they moved closer together. Both assays were conducted at 15°C, 20°C, 25°C and 30°C, with evaluation performed after 24 and 48 hours of incubation.

### 2.9. Characterization of Plant Growth-Promoting Traits and Stress Tolerance in the EcoBiome

*In vitro* evaluation of PGP traits for the EcoBiome was performed as previously described in the Section 2.3, with modifications. Briefly, the assay was conducted using individual isolates, pairwise combinations, and the complete EcoBiome, by inoculating 10 µL of each isolate as required.

The biofilm production capacity and water stress tolerance of the EcoBiome were evaluated following the protocols described by Chávez-Avila et al. (2023). For biofilm formation, isolates were cultured in Eppendorf tubes for six days without shaking. Subsequently, 0.3% crystal violet was added and incubated at room temperature for 15 minutes. The supernatant was discarded, and the tubes were washed with distilled water. The dye adhering to the tube walls was then solubilized with 1 mL of 100% ethanol, and absorbance was measured at 590 nm.

The water stress tolerance assay was performed on LB agar plates supplemented with 5% and 10% polyethylene glycol-8000 (PEG-8000) for 24 hours. In all assays, bacterial suspensions were adjusted to an optical density at 600 nm (OD_600_) of 0.1 and incubated at four temperatures:15°C, 20°C, 25°C and 30°C.

## 3. Results

### 3.1. Taxonomic identification and *In Vitro* Characterization of Plant Growth Promoting Traits

A total of 17 bacterial isolates were obtained from three Chilean desert soils: eight from the Atacama Desert, six from the Antarctic continent, and three from Las Cardas. In terms of taxonomic diversity, the Atacama Desert isolates were classified into seven different genera, while the Antarctic isolates belonged to only three genera. The three isolates from Las Cardas each represented a distinct genus.

The phylogenetic tree constructed using the 16S rRNA gene sequences (Figure 3) showed that the Antarctic Desert isolates formed a monophyletic clade with 91% bootstrap support. All members of this clade belonged to the order Micrococcales, with a well-supported internal group of closely related sequences from the genera *Arthrobacter* and *Pseudoarthrobacter*. A phylogenetically close isolate, *Streptomyces* sp. M1-A, was obtained from the Atacama Desert.

**Figure 3.**
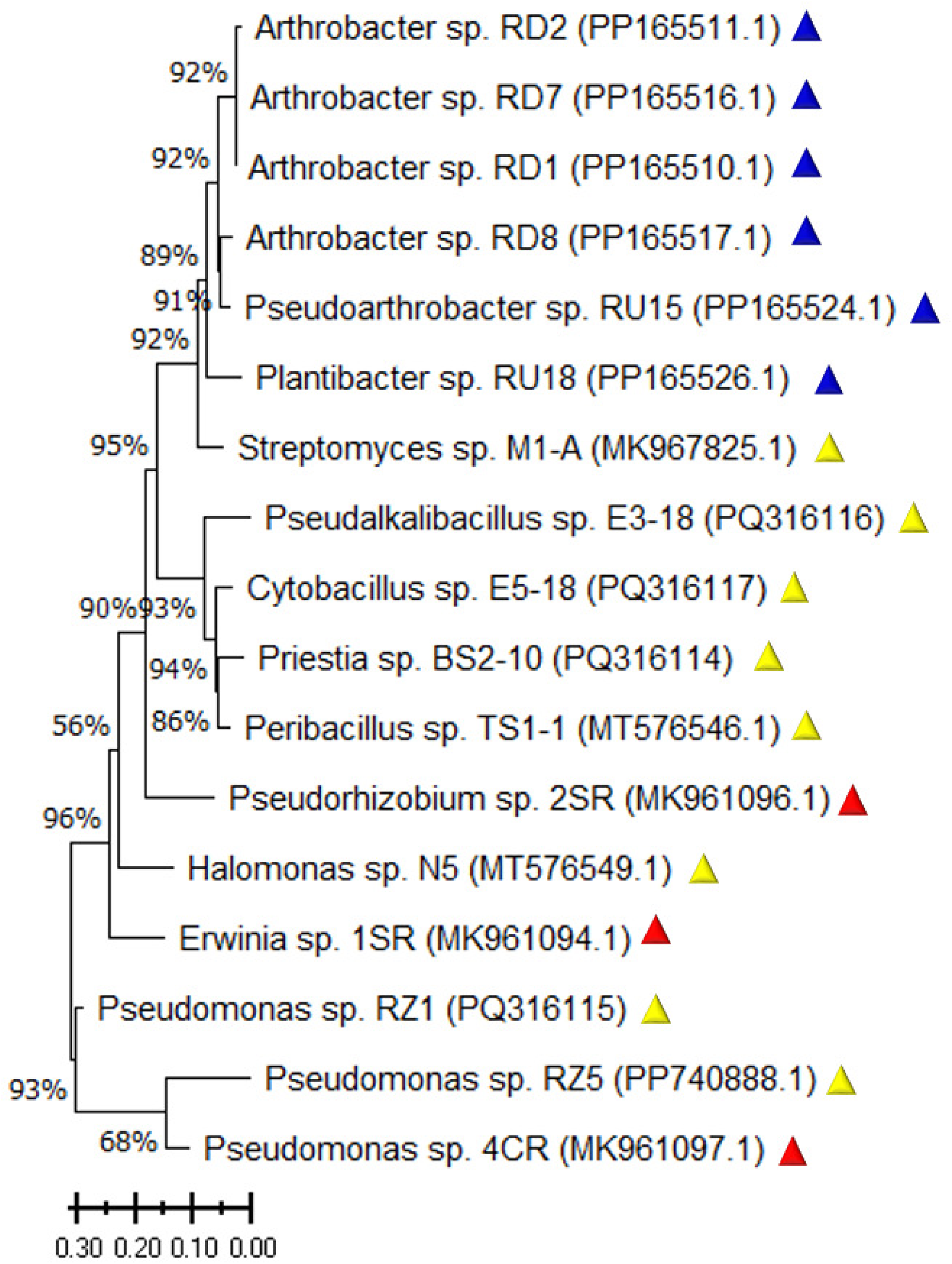
Phylogenetic tree based on 16S rRNA sequences of isolates from different desert environments. Colored triangles indicate the sampling sites: Antarctic Desert (blue), Atacama Desert (yellow), and Las Cardas (red).

Another observed clade is supported by a 93% bootstrap and includes the bacterial genera *Pseudalkalibacillus*, *Cytobacillus*, *Priestia* and *Peribacillus*, all included in the order *Bacillales* and isolated from the soil of the Atacama Desert.

Finally, another clade, with 90% bootstrap support, corresponded to the phylum Pseudomonadota and included three Atacama Desert and three Las Cardas isolates.

Subsequently, the PGP traits of each isolated bacteria were analyzed using specific culture media (Table 1), including siderophore production (CAS), phosphate solubilization (PKV), nitrification (NM), and indole-3-acetic acid (IAA), expressed in µg/mL. The results showed that 11 bacterial isolates exhibited a positive response for at least one PGP trait, specifically IAA production, with concentrations ranging from 0.99 µg/mL to 84.024 µg/mL. Notably, the *Plantibacter* sp. RU18 isolate (from the Antarctic Desert) demonstrated the highest IAA production. Several isolates exhibited multiple PGP traits. *Streptomyces* sp. M1-A and *Pseudorhizobium* sp. 2SR tested positive for both siderophore production (CAS) and IAA. *Pseudomonas* sp. 4CR showed positive activity in PKV, NM and IAA production. Meanwhile, *Pseudomonas* sp. RZ1, *Pseudomonas* sp. RZ5 and *Erwinia* sp. 1SR were positive for all four PGP traits: CAS, PKV, NM and IAA.

**Table 1.**
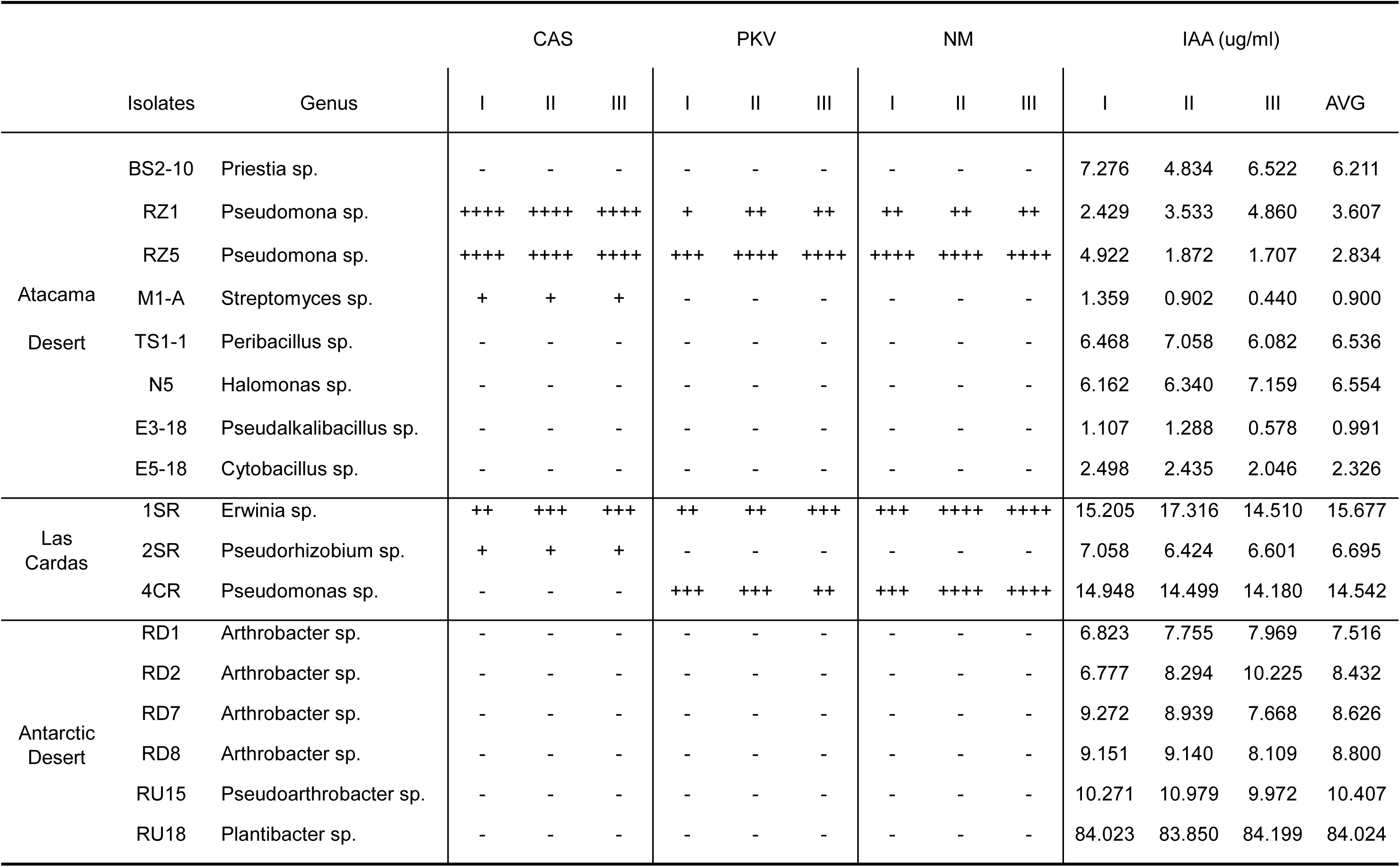
Summary of PGP traits characterization assays. The evaluated traits included siderophore production (CAS), phosphate solubilization (PKV), nitrification (NM), and indole-3- acetic acid (IAA) production. Columns I, II, and III represent triplicate measurements, while the fourth column in the IAA production assay indicates the average value (AVG). Halo size interpretation: (+) 1-2 mm; (++) 3-4 mm; (+++) 5-6 mm; (++++) 7-8 mm, measured from the edge of the bacterial colony to the edge of the halo formed.

### 3.2. Synthetic Community Assembly and Short-Term of Relative Abundance

To evaluate the changes in the relative abundance of the synthetic community in co-culture and assess the influence of temperature within an agronomically relevant range, the community was cultivated at 15°C, 20°C, 25°C, and 30°C for two consecutive subcultures (Figure 4).

**Figure 4.**
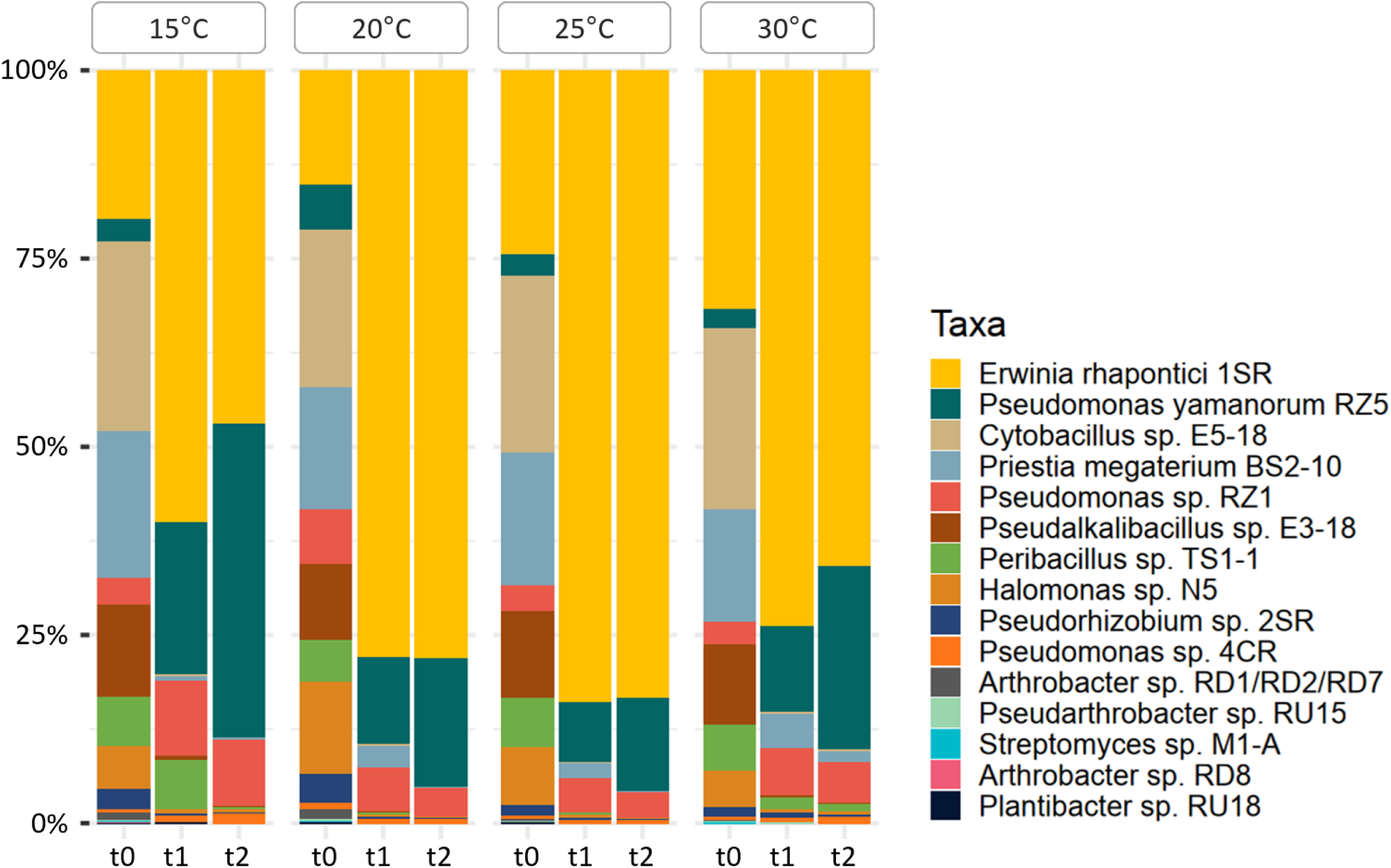
Relative abundance analysis of the synthetic community after two successive subcultures (t1 and t2) at four different agronomically relevant temperatures. Stacked bar plot of relative abundances based on 16S rRNA amplicon sequencing using Oxford Nanopore Technology.

At the beginning of the culture (t0), relative abundances were similar across all four temperatures for each isolate (Figure S1). However, after the first 24-hours subculture (t1), most isolates showed a decrease of at least 70%, or maintained their relative abundance, while approximately 30% of the isolates showed an increase. After the second subculture (t2), 90% of the isolates exhibited a further decrease in relative abundance, whereas the remaining 10% showed an increase in the culture medium.

Notably, following the second subculture (t2), two bacterial isolates became predominant in terms of relative abundance: *Erwinia* sp. 1SR, which increased between 20% and 60%, and *Pseudomonas* sp. RZ5, which increased between 10% and 40%. A slight increase was also observed in the relative abundance of *Pseudomonas* sp. RZ1.

### 3.3. Genome Sequencing and Phylogenomic Identification of Selected Isolates

To assemble the desert EcoBiome, we applied three selection criteria: (1) isolates must be representative of the native bacterial community of each desert, (2) isolates must exhibit PGP traits, and (3) isolates must show persistence or increased relative abundance across successive co- cultures. Based on these criteria, we selected one isolate from each desert: *Erwinia* sp. 1SR from Las Cardas and *Pseudomonas* sp. RZ5 from the Atacama Desert, both of which exhibited PGP traits and demonstrated the highest relative abundances during successive co-cultures. Additionally, *Plantibacter* sp. RU18 from the Antarctic continent was included due to its exceptionally high production of IAA, one of the most advantageous PGP traits observed in our strain collection.

To further investigate the metabolic capabilities of the selected isolates, we performed whole- genome sequencing. We obtained three high-quality genomes (Figure S2), and phylogenomic trees were constructed to assign genus- and species-level taxonomy (Figure S3). This genomic analysis allowed full taxonomic assignment for two of the three isolates: 1SR was identified as *Erwinia rhapontici* with 96% support, and RZ5 as *Pseudomonas yamanorum* with 100% support. For RU18, only the genus could be confirmed (*Plantibacter* sp.), with *Plantibacter flavus* as the closest species.

The genome assemblies (Table 2) revealed high completeness and low contamination: 99.86% completeness and 0.88% contamination for *Pseudomonas yamanorum* RZ5; 99.34% and 0.57% for *Erwinia rhapontici* 1SR; and 99.49% and 1.68% for *Plantibacter* sp. RU18. All genomes were fully circularized. In the cases of 1SR and RU18, two additional complete circular plasmids were also identified. The total assembly length was 7,257,956 bp for *Pseudomonas yamanorum* RZ5, with 6,600 predicted coding sequences (CDSs); *Erwinia rhapontici* 1SR had an assembly length of 5,439,041 bp with three contigs and 5,107 CDSs; and *Plantibacter* sp. RU18 had 4,710,248 bp and 4,463 CDSs.

**Table 2.** Molecular features obtained from genomic sequencing.

| Features | *Pseudomonas yamanorum* RZ5 | *Erwinia rhapontici* 1SR | *Plantibacter* sp. RU18 |
| --- | --- | --- | --- |
| ID | RZ5 | 1SR | RU18 |
| %Completeness | 99.86 | 99.34 | 99.49 |
| %Contamination | 0.88 | 0.57 | 1.68 |
| %Strain heterogeneity | 9.09 | 0 | 0 |
| N50 | 7,257,956 | 5,250,678 | 4,671,681 |
| Contigs | 1 | 3 | 3 |
| %GC | 60.3 | 54 | 67.4 |
| Total length | 7,257,956 bp | 5,439,041 bp | 4,710,248 bp |
| Predicted genes | 6740 | 5271 | 4522 |
| CDS | 6600 | 5107 | 4463 |
| pseudogenes | 44 | 42 | 26 |
| rRNA | 16 | 22 | 6 |
| tRNA | 70 | 84 | 50 |
| Genome Quality | High-quality draft | High-quality draft | High-quality draft |

### 3.4. Genome-Based Prediction of Secondary Metabolites and Metabolic Pathways

To further our characterization of PGP traits in the three selected isolates, we performed a screening for secondary metabolites and metabolic pathways using whole genome-based bioinformatics analyses.

A prediction of secondary metabolites was performed using antiSMASH (Table 3). Our results identified six different cluster types for *Erwinia rhapontici* 1SR, with two regions showing more than 50% similarity with known clusters. These corresponded to a NRPS (non-ribosomal peptide synthetase cluster) cluster (region 1. 3) with 61% of similarity to a siderophore production cluster, and region 1.2, which exhibited 94% similarity to an arylpolyene production cluster. On the other hand, *Pseudomonas yamanorum* RZ5 was predicted to have 15 gene clusters, of which three exhibited 50% or more similarity to known clusters: region 1.8 for pyocyanine (100%), region 1.10 for viscosin (50%), and region 1.12, showing 80% similarity to a pyoverdine cluster. Finally, in *Plantibacter* sp. RU18, we detected nine gene clusters, with two secondary metabolites showing similarities above 50%, corresponding to carotenoid (50%) and ε-Poly-L-lysine (100%).

**Table 3.**
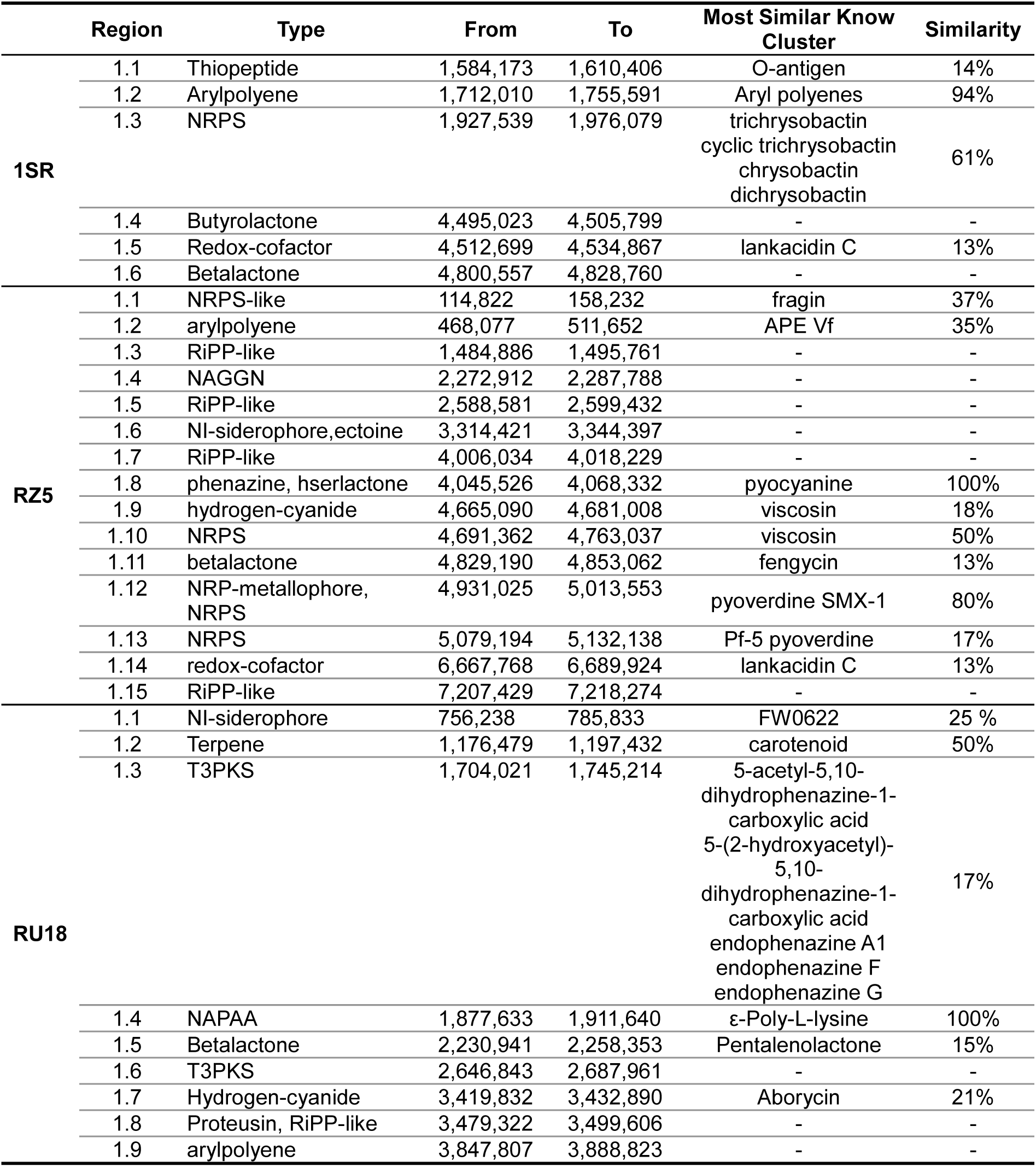
Secondary metabolites identified using the antiSMASH tool from genomic sequencing of the selected isolates *Erwinia rhapontici* 1SR, *Pseudomonas yamanorum* RZ5 and *Plantibacter* sp. RU18.

Additionally, we performed a genomic analysis using KofamKOALA to assign KEGG Orthologs (KOs) (Table S1). Two genes (*nirBD* and *nirD*) related to the dissimilatory nitrate reduction pathway were identified in *Plantibacter* sp. RU18. Furthermore, *Erwinia rhapontici* 1SR exhibited all screened genes associated with the phosphate uptake pathway (*pstS*, *pstA*, *pstC*, *pstB*, *phoA, phoR*, *phoU* and *phoB*), the iron uptake pathway (*fhuB*, *fhuC*, *fhuD*, *feoA* and *fur*), and the siderophore pathway (*fepB*, *fepC*, *fepD*, *fepG*). In contrast, these genes were only partially present in *Pseudomonas yamanorum* RZ5 and *Plantibacter* sp. RU18.

All isolates possessed the *efeB* gene, involved in the iron transport pathway, as well as some genes related to the IAA synthesis pathway (*mao, trpA*, *trpB*, *trpC* and *trpS*). Additionally, ACC deaminase (*acdS*) was absent only in *Erwinia rhapontici* 1SR.

We also analyzed metabolic pathways associated with biofilm formation, identifying 20 genes. The results indicated that *Pseudomonas yamanorum* RZ5 harbored all biofilm-related genes, while *Erwinia rhapontici* 1SR contained 13 genes, and *Plantibacter* sp. RU18 three. Furthermore, all isolates presented the murJ gene, associated with flagellar activity.

Finally, we analyzed genes associated with tolerance to abiotic stress conditions. In all three isolates, we detected genes related to heat stress tolerance (*smpB*, *dnaJ*, *dnaK*, *grpE*, *groES*, *clpB*, *clpX*, *htpX)* and genes associated with tolerance to drought and saline conditions (*nhaA*, *proB*, *proS*, *kdpA*, *kdpB*, *kdpF* and *kdoC*).

### 3.5. Growth kinetics and Competitive Interactions in the EcoBiome

To assess the effect of temperature on the growth kinetics of the selected isolates, growth curves were generated (Figure S4). The maximal optical density (OD_max_) and the doubling time (DT) were calculated (Table 4). The results indicated that the optimal temperature range for achieving the highest OD_max_ was between 15°C and 25°C. Additionally, a pronounced decrease in OD_max_ was observed for *Plantibacter* sp. RU18 at 30°C, whereas *Pseudomonas yamanorum* RZ5 and *Erwinia rhapontici* 1SR exhibited only slight decreases. Regarding the DT values, the lowest DT was recorded at 30°C. The second lowest DT occurred at 20°C, representing a balance between higher proliferation rates (lower DT) and a higher OD_max_ values for all three isolates, followed by 25°C and 15°C.

**Table 4.** Bacterial growth kinetics at different temperatures in selected isolates. Maximal Optical Density (OD_max_) and Doubling Time (DT) were calculated based on each growth curve.

|  | Temperature |  |  |  |  |  |  |  |
| --- | --- | --- | --- | --- | --- | --- | --- | --- |
|  | 15°C |  | 20°C |  | 25°C |  | 30°C |  |
| Isolate | ODmax | DT (h) | ODmax | DT (h) | ODmax | DT (h) | ODmax | DT (h) |
| RZ5 | 1.597 | 10.26 | 1.548 | 7.672 | 1.672 | 9.06 | 1.440 | 3.548 |
| 1SR | 1.378 | 10.81 | 1.448 | 7.931 | 1.298 | 8.516 | 1.147 | 4.417 |
| RU18 | 1.276 | 16.72 | 1.280 | 10.83 | 1.275 | 11.44 | 0.394 | 7.847 |

Additionally, proliferation dynamics were evaluated at 15°C, 20°C, 25°C and 30°C to assess bacterial competition or synergism in both pairwise and three-isolate (EcoBiome) cultures (Figure S5). The results showed no evidence of competition (inhibition) or attraction at any evaluated temperature, but rather complementary growth among the different combinations.

### 3.6. Functional Characterization of Plant Growth-Promoting Traits and Abiotic Stress Tolerance in the EcoBiome

To evaluate the contribution of each isolate to PGP traits within the EcoBiome, a characterization assay was performed at four incubation temperatures (15°, 20°, 25°, and 30°C) using individual isolates, in pairwise combinations, and as a three-strain consortium (EcoBiome) (Figure 5 and Figure S6).

**Figure 5.**
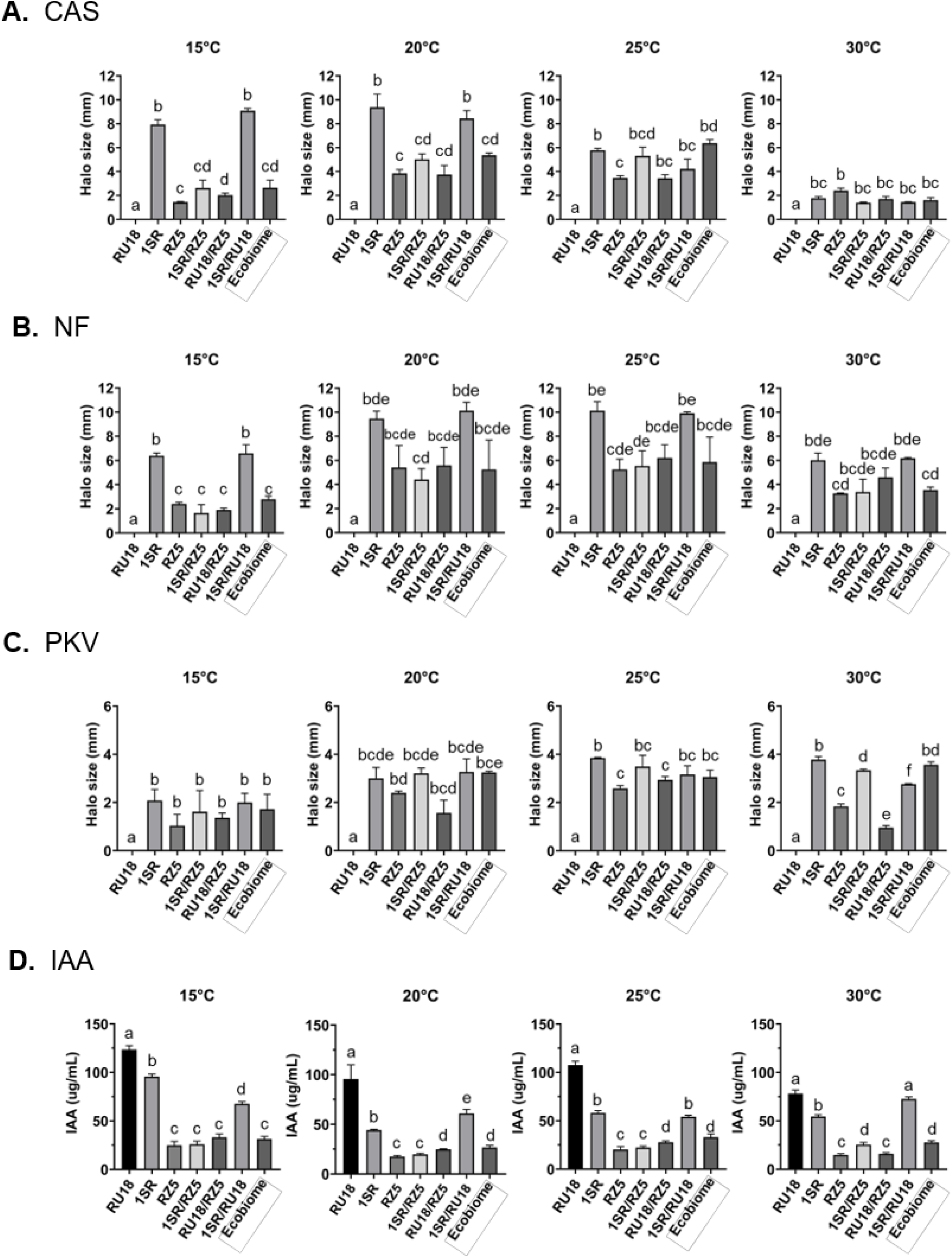
Characterization of plant growth-promoting (PGP) traits in *Erwinia rhapontici* 1SR, *Pseudomonas yamanorum* RZ5 and *Plantibacter sp.* RU18, evaluated individually, in pairwise combinations, and as the EcoBiome at four incubation temperatures. The *in vitro* assays included: (A) siderophore production on CAS agar, (B) nitrification process on NM agar, (C) phosphate solubilization on PKV agar, and (D) IAA production with Salkowski reagent.

The results reveled differences in siderophore production (CAS), nitrification (NM), phosphate solubilization (PKV), and indole acetic acid (IAA) production depending on the incubation temperature. Specifically, the highest siderophore production was observed at 20°C by *Erwinia rhapontici* 1SR, both individually and in combination with *Plantibacter* sp. RU18. In contrast, the EcoBiome showed no significant differences compared to *Pseudomonas yamanorum* RZ5 alone or its combinations with *Plantibacter* sp. RU18 or *Erwinia rhapontici* 1SR. A similar trend was observed at 15°C and 25°C, albeit with smaller halo size in all cases. At 30°C, the isolates, pairwise combinations, and the EcoBiome exhibited the lowest siderophore production, with no significant differences among them.

Regarding nitrification and phosphate solubilization values, the highest values were observed at 20°C and 25°C, with no significant differences between EcoBiome, *Erwinia rhapontici* 1SR, *Pseudomonas yamanorum* RZ5, and their combinations with *Plantibacter* sp. RU18. These values decreased at both 30°C and 15°C.

IAA production results indicated that *Plantibacter* sp. RU18 was the highest producer at all four tested temperatures, showing significant differences compared to the other isolates and their combinations. Meanwhile, the EcoBiome exhibited its highest IAA production at 25°C, with no significant differences compared to the combination of *Plantibacter* sp. RU18 and *Pseudomonas yamanorum* RZ5.

Additionally, two tests were performed to assess the adherence capacity and establishment of the EcoBiome: a biofilm formation assay (Figure 6), and assay evaluating proliferation under water deficit using polyethylene glycol (PEG) in the culture medium (Figure S7). The results showed that the EcoBiome formed significantly more biofilm at 25°C compared to each isolate or pairwise combination. At 15°C, 20°C, and 30°C, biofilm formation was lower, but the differences were not statistically significant compared to the bacterial pairs at each temperature.

**Figure 6.**
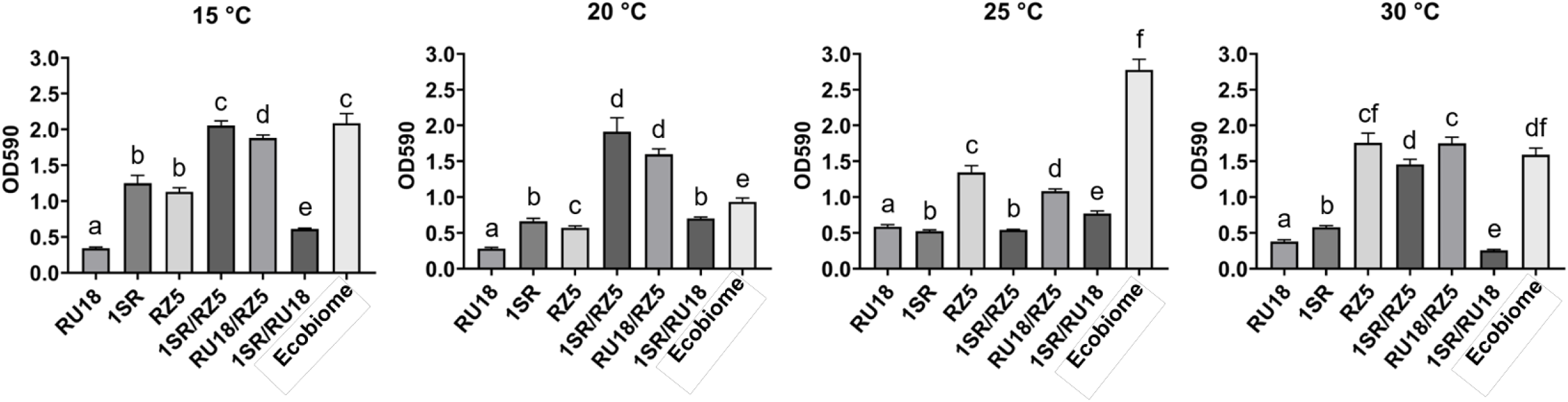
Biofilm formation assay of the EcoBiome and isolates. The assay was performed with *Erwinia rhapontici* 1SR, *Pseudomona yamanorum* RZ5, *Plantibacter* sp. RU18 individually, in pairwise combinations, and as the EcoBiome at four incubation temperatures.

On the other hand, *In Vitro* tolerance to water deficit revealed that all isolates tolerate both PEG 5% and PEG 10% conditions relative to the control (PEG 0%). The EcoBiome exhibited greater tolerance (measured as higher proliferation) at 25°C, followed by 20°C, 30°C, and 15°C for both PEG concentrations

## 4. Discussion

To develop a microbial community, using microorganisms from three desert environments with plant growth-promoting (PGP) properties, we integrated elements of both *top-down* and *bottom-up* design approaches. The *top-down* approach leverages environmental variables to select and direct microbial communities toward specific biological processes, while the *bottom-up* approach predicts how the metabolic attributes of individual microorganisms interact to achieve the desired function (Lawson et al., 2019). The combination of both strategies enabled us to construct an EcoBiome that preserves the adaptability of native desert communities, reduces the complexity of microbial interactions, and maintains the metabolic attributes necessary for the proposed function.

From the culturable fraction of each desert environment, we isolated diverse microorganisms using a nutrient-rich Luria-Bertani (LB) medium and a three-temperature culture scheme. Although the culturable fraction does not fully represent the native desert soil communities, this strategy, based on the top-down approach, simplified the system by reducing microbial diversity and addressing the challenges associated with managing complex microbial consortia (Mousa et al., 2024).

The analysis of PGP traits in the 17 isolates from the three deserts indicated that six exhibited at least one PGP trait. The evaluated traits included siderophore production, IAA generation, phosphate solubilization and nitrification (Table 1). Half of the isolates with PGP metabolic traits belonged to the genus *Pseudomonas*, consistent with several reports describing the PGP potential of *Pseudomonas* spp. (Gaete et al., 2022; Kour et al., 2020; Tian et al., 2023; Wei and Zhang, 2006). Additionally, the isolate from Las Cardas, identified as *Erwinia,* exhibited all the PGP traits tested. Previous investigations have described PGP capabilities in *Erwinia*, primarily in association with wheat (Egorshina et al., 2025) and rice plants (Jia et al., 2022). From Antarctic soils, the *Plantibacter* isolate showed the highest production of IAA in the entire collection, aligning with reports of *Plantibacter flavus* isolates from plant endospheres exhibiting PGP traits that benefited *Arabidopsis thaliana* by increasing biomass and root length (Mayer et al., 2019).

A genomic analysis to identify PGP traits was conducted through secondary metabolite screening using antiSMASH (Table 3), confirming the biochemical assays. Both *Erwinia rhapontici* 1SR and *Pseudomonas yamanorum* RZ5 harbored siderophore-related gene clusters with >60% similarity to reference clusters. The significance of siderophores in Erwinia species has been previously reported, particularly for *Erwinia amylovora* under iron-limitation conditions in apple flowers (Müller et al., 2022). Similarly, *Erwinia* sp. QZ-E9, isolated from *Sesuvium portulacastrum,* has been described as a key PGP bacterium (Cen et al., 2024). Siderophore production by *Pseudomonas yamanorum* was also recently reported by Licciardello et al. (2025), who isolated this bacterium from *Colobanthus quitensis* in Antarctic environments.

Moreover, the *Pseudomonas yamanorum* RZ5 isolate contained a pyocyanin biosynthetic gene cluster, with 100% similarity to characterized clusters. Pyocyanin, predominantly associated with *Pseudomonas aeruginosa,* has been implicated in biofilm formation (Das et al., 2015), potentially by enhancing electron transport within biofilms or modifying their viscosity (Das et al., 2015; Shouman et al., 2023). Pyocyanin, also participates in iron metabolism (Jayaseelan et al., 2014). Additionally, we detected a pyoverdine cluster (80% similarity), a well-known siderophore crucial for iron acquisition in *Pseudomonas fluorescens*, which indirectly protects plants from phytopathogens (Guo et al., 2021; Visca et al., 2007). Iron limitation, notably in pyoverdine-deficient mutants, has been linked to altered bacterial motility and suppressed biofilm formation (Guan et al., 2024).

Another noteworthy cluster in *Pseudomonas yamanorum* RZ5 was that of viscosin, a non-ribosomal peptide synthetase (NRPS) product. Viscosin has been implicated in microbial community establishment at the root-soil interface in germinating seeds, improving colonization by wild-type *Pseudomonas fluorescens* strains compared to viscosin-deficient mutants (Guan et al., 2024).

The secondary metabolite analysis performed on the *Erwinia rhapontici* 1SR isolate highlighted the presence of arylpolyenes, with 94% similarity to a known cluster. Arylpolyenes (APEs) are pigments produced by several bacteria genera that have been recently studied for their protective role against photooxidative damage and lipid peroxidation (Pailliè-Jiménez et al., 2024), similar to the effect of carotenoids (Schöner et al., 2016). Another identified secondary metabolite, with a 61% similarity to a known cluster, corresponded to siderophores, including chrysobactin, dichrysobactin, trichrysobactin, and cyclic trichrysobactin, which capture iron (III) from the environment and deliver it to cells (Sandy and Butler, 2011). Previous studies have reported the production of the siderophore catechol chrysobactin (Rauscher et al., 2002) in *Erwinia chrysanthemi*, and we observed that this PGP trait was maintained in the isolates when tested individually, in pairwise combinations and as part of the EcoBiome.

In *Plantibacter* sp. RU18, we detected with 100% similarity the presence of ε-poly-L-lysine, a linear homopoly (amino acid) whose characteristics as a biopolymer depend on its molecular weight and the producing species. This compound has been primarily used in industrial applications as a food preservative due to its antimicrobial activity against Gram-positive and Gram-negative bacteria at concentrations of 1-8 μg/ml (Shima et al., 1984), with a mechanism of action based on disruption of membrane integrity and permeability (Wang et al., 2021; Ye et al., 2013).

In addition to the genomic search for secondary metabolites, genes associated with PGP properties were also identified using KofamKOALA to assign KEGG Orthologs (KOs) (Table S1). Genes related to the phosphate solubilization, siderophore production, and nitrate dissimilatory reduction were identified in the genome of the three isolates selected to comprise the EcoBiome. The presence of genes associated with PGP traits correlated with the biochemical assay results for *Erwinia rhapontici* 1SR and *Pseudomonas yamanorum* RZ5. It is noteworthy that Erwinia rhapontici 1SR carried genes related to phosphate uptake, such as pstSACB and phoABRU, in addition to trpAB genes, which encode the alpha and beta subunits of tryptophan synthase (Li et al., 2021). However, discrepancies between genomic and phenotypic profiles emerged in *Plantibacter* sp. RU18. Although genes associated with nitrogen metabolism, phosphate solubilization and siderophore production were identified, these traits were not detected in the plate assays, suggesting absence of complete metabolic pathways or misclassification due to low identity of the annotated genes.

It has been proposed that commonly studied PGP traits, such as IAA production, phosphate solubilization (PKV), siderophore activity (CAS) and biofilm formation, correlate with ecological role of the isolates in the bacterial community and their host colonization efficiency (de Souza et al., 2019). Additionally, various bacterial species can stimulate plant growth both by inducing host metabolite production and by directly synthesizing beneficial compounds (Bal et al., 2013; Bastián et al., 1998; Wani and Khan, 2010).

Phenotypic results showed variation in the production of siderophore, nitrification and phosphate solubilization depending on temperature and microbial combinations. Siderophore production was the most sensitive to temperature changes, with a significant increase in halo size at 25°C, but a drastic reduction at 30°C. This behavior aligns with studies emphasizing the influence of environmental factors on siderophore production, with variations being species-dependent (Chaudhary et al., 2017). In our study, Erwinia rhapontici 1SR appeared to be the primary contributor to siderophore production in the EcoBiome, although its sensitivity to temperatures above 25°C suggests potential limitations under warmer conditions.

The production of indole-3-acetic acid (IAA) was notable in *Plantibacter* sp. RU18, which exhibited significantly higher levels of IAA production than the other two isolates and their combinations. This result aligns with previous reports of high IAA production in *Plantibacter* species, reaching values above 85 µg/mL (Wang et al., 2020). Other *Plantibacter* species with PGP traits have been reported, contributing to improved growth in *Arabidopsis thaliana* and carrying genes associated with auxin and cytokinin synthesis (Mayer et al., 2019), suggesting that this genus could play a key role in regulating plant growth.

Additionally, we evaluated biofilm formation individually, in pairwise combinations, and in the EcoBiome. Our results showed that the EcoBiome enhances biofilm production, consistent with previous findings where bacterial consortia produced more biofilm than individual isolates (Tian et al., 2023). Biofilm are multicellular communities embedded in extracellular matrices that protect microorganisms and enhance survival (Elumalai et al., 2024). These structures facilitate microbial coexistence in natural environments, promoting community stability.

The coexistence of these three isolates without significant antagonism highlights their potential for use in microbial inoculants for biotechnological applications in agriculture. The formation of stable, multispecies biofilms is essential for niche colonization and efficient plant colonization (Kapoore et al., 2022). Within biofilms, the extracellular polymeric substance (EPS) matrix, composed of nucleic acids, polysaccharides, lipids, and other biomolecules, plays a critical role in structural integrity (Elumalai et al., 2024). However, EPS production is influenced by environmental factors such as temperature, which may affect biofilm stability in certain ecological niches (Naylor and Coleman-Derr, 2018).

## 5. Conclusion

In this study, we successfully designed and characterized a synthetic microbial community (EcoBiome) composed of three bacterial isolates, *Erwinia rhapontici* 1SR, *Pseudomonas yamanorum* RZ5, and *Plantibacter* sp. RU18, originating from distinct desert environments. By integrating top-down and bottom-up design strategies, we selected strains that demonstrated complementary plant growth-promoting (PGP) traits, including phosphate solubilization, siderophore and IAA production, nitrogen metabolism, and biofilm formation.

Our results revealed that these traits were influenced by incubation temperature and microbial interaction, with the EcoBiome outperforming individual isolates in certain functional attributes, such as biofilm production. Genomic analysis confirmed the presence of biosynthetic gene clusters associated with secondary metabolites and PGP pathways, which correlated with the phenotypic assays, especially for E. rhapontici and P. yamanorum. Despite Plantibacter sp. RU18 showing limited functional expression in vitro for some traits, its strong IAA production and metabolic potential support its inclusion in the consortium.

Additionally, the EcoBiome demonstrated the ability to tolerate water deficit conditions, maintaining its growth under reduced water availability. This feature, combined with biofilm formation capacity, suggests potential ecological resilience in arid soils where water availability is limited.

The compatibility, functional complementarity, and ecological resilience of the EcoBiome suggest its potential application as a microbial inoculant in arid and semi-arid agroecosystems. Further studies under greenhouse and field conditions are needed to evaluate its effectiveness in promoting plant growth and enhancing crop productivity under environmental stress.

## Acknowledgments

This work was supported by Millennium Science Initiative Program Center of Genome Regulation ICN2021_044, ANID FONDECYT grants 1241424 to MG and 3250795 to AG.

## References

1. Akhtyamova, Z., Martynenko, E., Arkhipova, T., Seldimirova, O., Galin, I., Belimov, A., Vysotskaya, L., Kudoyarova, G., 2023. Influence of Plant Growth-Promoting Rhizobacteria on the Formation of Apoplastic Barriers and Uptake of Water and Potassium by Wheat Plants. Microorganisms 11, 1227. 10.3390/microorganisms11051227

2. Alsharif, W., Saad, M.M., Hirt, H., 2020. Desert Microbes for Boosting Sustainable Agriculture in Extreme Environments. Front. Microbiol. 11. 10.3389/fmicb.2020.01666

3. Aramaki, T., Blanc-Mathieu, R., Endo, H., Ohkubo, K., Kanehisa, M., Goto, S., Ogata, H., 2020. KofamKOALA: KEGG Ortholog assignment based on profile HMM and adaptive score threshold. Bioinformatics 36, 2251–2252. 10.1093/bioinformatics/btz859

4. Backer, R., Rokem, J.S., Ilangumaran, G., Lamont, J., Praslickova, D., Ricci, E., Subramanian, S., Smith, D.L., 2018. Plant Growth-Promoting Rhizobacteria: Context, Mechanisms of Action, and Roadmap to Commercialization of Biostimulants for Sustainable Agriculture. Front Plant Sci 9, 1473. 10.3389/fpls.2018.01473

5. Bal, H.B., Das, S., Dangar, T.K., Adhya, T.K., 2013. ACC deaminase and IAA producing growth promoting bacteria from the rhizosphere soil of tropical rice plants. Journal of Basic Microbiology 53, 972–984. 10.1002/jobm.201200445

6. Bardgett, R.D., Caruso, T., 2020. Soil microbial community responses to climate extremes: resistance, resilience and transitions to alternative states. Philosophical Transactions of the Royal Society B: Biological Sciences 375, 20190112. 10.1098/rstb.2019.0112

7. Bastián, F., Cohen, A., Piccoli, P., Luna, V., Bottini*, R., Baraldi, R., Bottini, R., 1998. Production of indole-3-acetic acid and gibberellins A1 and A3 by Acetobacter diazotrophicus and Herbaspirillum seropedicae in chemically-defined culture media. Plant Growth Regulation 24, 7–11. 10.1023/A:1005964031159

8. Berendsen, R.L., Vismans, G., Yu, K., Song, Y., De Jonge, R., Burgman, W.P., Burmølle, M., Herschend, J., Bakker, P.A.H.M., Pieterse, C.M.J., 2018. Disease-induced assemblage of a plant- beneficial bacterial consortium. The ISME Journal 12, 1496–1507. 10.1038/s41396-018-0093-1

9. Blin, K., Shaw, S., Augustijn, H.E., Reitz, Z.L., Biermann, F., Alanjary, M., Fetter, A., Terlouw, B.R., Metcalf, W.W., Helfrich, E.J.N., van Wezel, G.P., Medema, M.H., Weber, T., 2023. antiSMASH 7.0: new and improved predictions for detection, regulation, chemical structures and visualisation. Nucleic Acids Research 51, W46–W50. 10.1093/nar/gkad344

10. Bokszczanin, K.Ł., Wrona, D., Przybyłko, S., 2021. The Effect of Microbial Inoculation under Various Nitrogen Regimes on the Uptake of Nutrients by Apple Trees. Agronomy 11, 2348. 10.3390/agronomy11112348

11. Cavazos, B.R., Bohner, T.F., Donald, M.L., Sneck, M.E., Shadow, A., Omacini, M., Rudgers, J.A., Miller, T.E.X., 2018. Testing the roles of vertical transmission and drought stress in the prevalence of heritable fungal endophytes in annual grass populations. New Phytologist 219, 1075–1084. 10.1111/nph.15215

12. Cen, X., Li, H., Zhang, Y., Huang, L., Luo, Y., 2024. Isolation and Plant Growth Promotion Effect of Endophytic Siderophore-Producing Bacteria: A Study on Halophyte Sesuvium portulacastrum. Plants 13, 2703. 10.3390/plants13192703

13. Chaudhary, D.Y., Gosavi, P., Durve-Gupta, A., 2017. Isolation and application of siderophore producing bacteria. International Journal of Applied Research 3, 246–250.

14. Chaumeil, P.-A., Mussig, A.J., Hugenholtz, P., Parks, D.H., 2020. GTDB-Tk: a toolkit to classify genomes with the Genome Taxonomy Database. Bioinformatics 36, 1925–1927. 10.1093/bioinformatics/btz848

15. Chávez-Avila, S., Valencia-Marin, M.F., Guzmán-Guzmán, P., Kumar, A., Babalola, O.O., Orozco-Mosqueda, M.D.C., De Los Santos-Villalobos, S., Santoyo, G., 2023. Deciphering the antifungal and plant growth-stimulating traits of the stress-tolerant Streptomyces achromogenes subsp. achromogenes strain UMAF16, a bacterium isolated from soils affected by underground fires. Biocatalysis and Agricultural Biotechnology 53, 102859. 10.1016/j.bcab.2023.102859

16. Chen, H., Costanza, R., 2024. Valuation and management of desert ecosystems and their services. Ecosystem Services 66, 101607. 10.1016/j.ecoser.2024.101607

17. Cheng, X., Cheng, H., Sun, G., Shi, X., 2017. Study of Changing Features of Precipitation from 1900-2010 Years in Africa-Asia Arid and Semi-Arid Area. Journal of Geoscience and Environment Protection 05, 62. 10.4236/gep.2017.51004

18. Cherif, H., Marasco, R., Rolli, E., Ferjani, R., Fusi, M., Soussi, A., Mapelli, F., Blilou, I., Borin, S., Boudabous, A., Cherif, A., Daffonchio, D., Ouzari, H., 2015. Oasis desert farming selects environment-specific date palm root endophytic communities and cultivable bacteria that promote resistance to drought. Environ Microbiol Rep 7, 668–678. 10.1111/1758-2229.12304

19. Curry, K.D., Wang, Q., Nute, M.G., Tyshaieva, A., Reeves, E., Soriano, S., Wu, Q., Graeber, E., Finzer, P., Mendling, W., Savidge, T., Villapol, S., Dilthey, A., Treangen, T.J., 2022. Emu: species-level microbial community profiling of full-length 16S rRNA Oxford Nanopore sequencing data. Nat Methods 19, 845–853. 10.1038/s41592-022-01520-4

20. Das, T., Kutty, S.K., Tavallaie, R., Ibugo, A.I., Panchompoo, J., Sehar, S., Aldous, L., Yeung, A.W.S., Thomas, S.R., Kumar, N., Gooding, J.J., Manefield, M., 2015. Phenazine virulence factor binding to extracellular DNA is important for Pseudomonas aeruginosa biofilm formation. Sci Rep 5, 8398. 10.1038/srep08398

21. de Souza, R.S.C., Armanhi, J.S.L., Damasceno, N. de B., Imperial, J., Arruda, P., 2019. Genome Sequences of a Plant Beneficial Synthetic Bacterial Community Reveal Genetic Features for Successful Plant Colonization. Front. Microbiol. 10. 10.3389/fmicb.2019.01779

22. Devi, R., Kaur, T., Kour, D., Yadav, A.N., 2022. Microbial consortium of mineral solubilizing and nitrogen fixing bacteria for plant growth promotion of amaranth (*Amaranthus hypochondrius* L.). Biocatalysis and Agricultural Biotechnology 43, 102404. 10.1016/j.bcab.2022.102404

23. Dhull, S., Gera, R., Sheoran, H.S., Kakar, R., 2018. Phosphate Solubilization Activity of Rhizobial Strains Isolated From Root Nodule of Cluster Bean Plant Native to Indian Soils. Int.J.Curr.Microbiol.App.Sci 7, 255–266. 10.20546/ijcmas.2018.704.029

24. Diniz, F.V., Scherwinski-Pereira, J.E., Costa, F.H.S., Carvalho, C.M., 2025. Effects on plant physiology in response to inoculation of growth-promoting bacteria: systematic review. Braz. J. Biol. 85, e287279. 10.1590/1519-6984.287279

25. Egorshina, A., Lukyantsev, M., Golubev, S., Boulygina, E., Khilyas, I., Muratova, A., 2025. Erwinia plantamica sp. nov., a Non-Phytopathogenic Bacterium Isolated from the Seedlings of Spring Wheat (Triticum aestivum L.). Microorganisms 13, 474. 10.3390/microorganisms13030474

26. Elumalai, P., Gao, X., Cui, J., Kumar, A.S., Dhandapani, P., Parthipan, P., Karthikeyan, O.P., Theerthagiri, J., Kheawhom, S., Choi, M.Y., 2024. Biofilm formation, occurrence, microbial communication, impact and characterization methods in natural and anthropic systems: a review. Environ Chem Lett 22, 1297–1326. 10.1007/s10311-024-01715-5

27. Farris, J.S., 1972. Estimating Phylogenetic Trees from Distance Matrices. The American Naturalist 106, 645–668. 10.1086/282802

28. Fierer, N., Lauber, C.L., Ramirez, K.S., Zaneveld, J., Bradford, M.A., Knight, R., 2012. Comparative metagenomic, phylogenetic and physiological analyses of soil microbial communities across nitrogen gradients. The ISME Journal 6, 1007–1017. 10.1038/ismej.2011.159

29. Gaete, A., Andreani-Gerard, C., Maldonado, J.E., Muñoz-Torres, P.A., Sepúlveda-Chavera, G.F., González, M., 2022. Bioprospecting of Plant Growth-Promoting Traits of Pseudomonas sp. Strain C3 Isolated from the Atacama Desert: Molecular and Culture-Based Analysis. Diversity 14, 388. 10.3390/d14050388

30. Gaete, A., Mandakovic, D., González, M., 2020. Isolation and Identification of Soil Bacteria from Extreme Environments of Chile and Their Plant Beneficial Characteristics. Microorganisms 8, 1213. 10.3390/microorganisms8081213

31. Gaete, A., Pulgar, R., Hodar, C., Maldonado, J., Pavez, L., Zamorano, D., Pastenes, C., González, M., Franck, N., Mandakovic, D., 2021. Tomato Cultivars With Variable Tolerances to Water Deficit Differentially Modulate the Composition and Interaction Patterns of Their Rhizosphere Microbial Communities. Front. Plant Sci. 12. 10.3389/fpls.2021.688533

32. Gao, Q., Garcia-Pichel, F., 2011. Microbial ultraviolet sunscreens. Nat Rev Microbiol 9, 791–802. 10.1038/nrmicro2649

33. Ge, Y., Abdulkreem AL-Huqail, A., Zhou, Z., Ali, E.F., Ghoneim, A.M., Eissa, M., El-Sharkawy, M.S., Ding, Z., 2022. Plant Growth Stimulating Bacteria and Filter Mud Cake Enhance Soil Quality and Productivity of Mango (Mangifera indica L.). J Soil Sci Plant Nutr 22, 3068–3080. 10.1007/s42729-022-00868-y

34. Gopalakrishnan, V., Burdman, S., Jurkevitch, E., Helman, Y., 2022. From the Lab to the Field: Combined Application of Plant-Growth-Promoting Bacteria for Mitigation of Salinity Stress in Melon Plants. Agronomy 12, 408. 10.3390/agronomy12020408

35. Guan, Y., Bak, F., Hennessy, R.C., Horn Herms, C., Elberg, C.L., Dresbøll, D.B., Winding, A., Sapkota, R., Nicolaisen, M.H., 2024. The potential of Pseudomonas fluorescens SBW25 to produce viscosin enhances wheat root colonization and shapes root-associated microbial communities in a plant genotype-dependent manner in soil systems. mSphere 9, e0029424. 10.1128/msphere.00294-24

36. Guo, W., Li, F., Xia, J., Wang, W., 2021. Complete genome sequence of a marine-derived bacterium Pseudomonas sp. SXM-1 and characterization of its siderophore through antiSMASH analysis and with mass spectroscopic method. Mar Genomics 55, 100802. 10.1016/j.margen.2020.100802

37. He, T., Li, Z., Sun, Q., Xu, Y., Ye, Q., 2016. Heterotrophic nitrification and aerobic denitrification by *Pseudomonas tolaasii* Y-11 without nitrite accumulation during nitrogen conversion. Bioresource Technology 200, 493–499. 10.1016/j.biortech.2015.10.064

38. Hunt, M., Silva, N.D., Otto, T.D., Parkhill, J., Keane, J.A., Harris, S.R., 2015. Circlator: automated circularization of genome assemblies using long sequencing reads. Genome Biol 16, 294. 10.1186/s13059-015-0849-0

39. Jayaseelan, S., Ramaswamy, D., Dharmaraj, S., 2014. Pyocyanin: production, applications, challenges and new insights. World J Microbiol Biotechnol 30, 1159–1168. 10.1007/s11274-013-1552-5

40. Jia, H., Xi, Z., Ma, J., Li, Y., Hao, C., Lu, M., Zhang, Z.-Z., Deng, W.-W., 2022. Endophytic bacteria from the leaves of two types of albino tea plants, indicating the plant growth promoting properties. Plant Growth Regul 96, 331–343. 10.1007/s10725-021-00779-5

41. Jooste, M., Roets, F., Midgley, G.F., Oberlander, K.C., Dreyer, L.L., 2019. Nitrogen-fixing bacteria and Oxalis – evidence for a vertically inherited bacterial symbiosis. BMC Plant Biol 19, 441. 10.1186/s12870-019-2049-7

42. Kalvari, I., Nawrocki, E.P., Ontiveros-Palacios, N., Argasinska, J., Lamkiewicz, K., Marz, M., Griffiths-Jones, S., Toffano-Nioche, C., Gautheret, D., Weinberg, Z., Rivas, E., Eddy, S.R., Finn, R.D., Bateman, A., Petrov, A.I., 2020. Rfam 14: expanded coverage of metagenomic, viral and microRNA families. Nucleic Acids Res 49, D192–D200. 10.1093/nar/gkaa1047

43. Kapoore, R.V., Padmaperuma, G., Maneein, S., Vaidyanathan, S., 2022. Co-culturing microbial consortia: approaches for applications in biomanufacturing and bioprocessing. Critical Reviews in Biotechnology 42, 46–72. 10.1080/07388551.2021.1921691

44. Karlidag, H., Esitken, A., Turan, M., Sahin, F., 2007. Effects of root inoculation of plant growth promoting rhizobacteria (PGPR) on yield, growth and nutrient element contents of leaves of apple. Scientia Horticulturae 114, 16–20. 10.1016/j.scienta.2007.04.013

45. Katsenios, N., Andreou, V., Sparangis, P., Djordjevic, N., Giannoglou, M., Chanioti, S., Stergiou, P., Xanthou, M.-Z., Kakabouki, I., Vlachakis, D., Djordjevic, S., Katsaros, G., Efthimiadou, A., 2021. Evaluation of Plant Growth Promoting Bacteria Strains on Growth, Yield and Quality of Industrial Tomato. Microorganisms 9, 2099. 10.3390/microorganisms9102099

46. Khan, M.Y., Nadeem, S.M., Sohaib, M., Waqas, M.R., Alotaibi, F., Ali, L., Zahir, Z.A., Al-Barakah, F.N.I., 2022. Potential of plant growth promoting bacterial consortium for improving the growth and yield of wheat under saline conditions. Front. Microbiol. 13. 10.3389/fmicb.2022.958522

47. Kochhar, N., I․k, K., Shrivastava, S., Ghosh, A., Rawat, V.S., Sodhi, K.K., Kumar, M., 2022. Perspectives on the microorganism of extreme environments and their applications. Current Research in Microbial Sciences 3, 100134. 10.1016/j.crmicr.2022.100134

48. Kolmogorov, M., Yuan, J., Lin, Y., Pevzner, P.A., 2019. Assembly of long, error-prone reads using repeat graphs. Nat Biotechnol 37, 540–546. 10.1038/s41587-019-0072-8

49. Kour, D., Rana, K.L., Sheikh, I., Kumar, V., Yadav, A.N., Dhaliwal, H.S., Saxena, A.K., 2020. Alleviation of Drought Stress and Plant Growth Promotion by Pseudomonas libanensis EU-LWNA- 33, a Drought-Adaptive Phosphorus-Solubilizing Bacterium. Proc. Natl. Acad. Sci., India, Sect. B Biol. Sci. 90, 785–795. 10.1007/s40011-019-01151-4

50. Krzywinski, M., Schein, J., Birol, I., Connors, J., Gascoyne, R., Horsman, D., Jones, S.J., Marra, M.A., 2009. Circos: an information aesthetic for comparative genomics. Genome Res 19, 1639– 1645. 10.1101/gr.092759.109

51. Kumar, A., Verma, J.P., 2018. Does plant—Microbe interaction confer stress tolerance in plants: A review? Microbiological Research 207, 41–52. 10.1016/j.micres.2017.11.004

52. Larsson, A., 2014. AliView: a fast and lightweight alignment viewer and editor for large datasets. Bioinformatics 30, 3276–3278. 10.1093/bioinformatics/btu531

53. Lawson, C.E., Harcombe, W.R., Hatzenpichler, R., Lindemann, S.R., Löffler, F.E., O’Malley, M.A., García Martín, H., Pfleger, B.F., Raskin, L., Venturelli, O.S., Weissbrodt, D.G., Noguera, D.R., McMahon, K.D., 2019. Common principles and best practices for engineering microbiomes. Nat Rev Microbiol 17, 725–741. 10.1038/s41579-019-0255-9

54. Lefort, V., Desper, R., Gascuel, O., 2015. FastME 2.0: A Comprehensive, Accurate, and Fast Distance-Based Phylogeny Inference Program. Mol Biol Evol 32, 2798–2800. 10.1093/molbev/msv150

55. Leger, A., Leonardi, T., 2019. pycoQC, interactive quality control for Oxford Nanopore Sequencing. Journal of Open Source Software 4, 1236. 10.21105/joss.01236

56. Li, H., 2018. Minimap2: pairwise alignment for nucleotide sequences. Bioinformatics 34, 3094– 3100. 10.1093/bioinformatics/bty191

57. Li, H., 2016. Minimap and miniasm: fast mapping and de novo assembly for noisy long sequences. Bioinformatics 32, 2103–2110. 10.1093/bioinformatics/btw152

58. Li, T., Mann, R., Kaur, J., Spangenberg, G., Sawbridge, T., 2021. Transcriptome Analyses of Barley Roots Inoculated with Novel Paenibacillus sp. and Erwinia gerundensis Strains Reveal Beneficial Early-Stage Plant–Bacteria Interactions. Plants 10, 1802. 10.3390/plants10091802

59. Licciardello, G., Antonielli, L., Sicher, C., Larini, I., Perazzolli, M., 2025. Two Antarctic endophytic bacteria of Colobanthus quitensis show functional and genomic characteristics potentially responsible for plant growth promotion and cold tolerance. Polar Biol 48, 42. 10.1007/s00300-025-03367-9

60. Mandakovic, D., Aguado-Norese, C., García-Jiménez, B., Hodar, C., Maldonado, J.E., Gaete, A., Latorre, M., Wilkinson, M.D., Gutiérrez, R.A., Cavieres, L.A., Medina, J., Cambiazo, V., Gonzalez, M., 2023. Testing the stress gradient hypothesis in soil bacterial communities associated with vegetation belts in the Andean Atacama Desert. Environmental Microbiome 18, 24. 10.1186/s40793-023-00486-w

61. Mandakovic, D., Rojas, C., Maldonado, J., Latorre, M., Travisany, D., Delage, E., Bihouée, A., Jean, G., Díaz, F.P., Fernández-Gómez, B., Cabrera, P., Gaete, A., Latorre, C., Gutiérrez, R.A., Maass, A., Cambiazo, V., Navarrete, S.A., Eveillard, D., González, M., 2018. Structure and co-occurrence patterns in microbial communities under acute environmental stress reveal ecological factors fostering resilience. Sci Rep 8, 5875. 10.1038/s41598-018-23931-0

62. Marasco, R., Mosqueira, M.J., Cherif, A., Daffonchio, D., 2022. Diversity and Plant Growth- Promoting Properties of Microbiomes Associated with Plants in Desert Soils, in: Ramond, J.-B., Cowan, D.A. (Eds.), Microbiology of Hot Deserts. Springer International Publishing, Cham, pp. 205–233. 10.1007/978-3-030-98415-1_8

63. Mayer, E., Dörr de Quadros, P., Fulthorpe, R., 2019. Plantibacter flavus, Curtobacterium herbarum, Paenibacillus taichungensis, and Rhizobium selenitireducens Endophytes Provide Host-Specific Growth Promotion of Arabidopsis thaliana, Basil, Lettuce, and Bok Choy Plants. Appl Environ Microbiol 85, e00383–19. 10.1128/AEM.00383-19

64. Mehrabi, Z., McMillan, V.E., Clark, I.M., Canning, G., Hammond-Kosack, K.E., Preston, G., Hirsch, P.R., Mauchline, T.H., 2016. Pseudomonas spp. diversity is negatively associated with suppression of the wheat take-all pathogen. Sci Rep 6, 29905. 10.1038/srep29905

65. Meier-Kolthoff, J.P., Carbasse, J.S., Peinado-Olarte, R.L., Göker, M., 2022. TYGS and LPSN: a database tandem for fast and reliable genome-based classification and nomenclature of prokaryotes. Nucleic Acids Research 50, D801. 10.1093/nar/gkab902

66. Meier-Kolthoff, J.P., Göker, M., 2019. TYGS is an automated high-throughput platform for state-of- the-art genome-based taxonomy. Nat Commun 10, 2182. 10.1038/s41467-019-10210-3

67. Mirskaya, G.V., Khomyakov, Y.V., Rushina, N.A., Vertebny, V.E., Chizhevskaya, E.P., Chebotar, V.K., Chesnokov, Y.V., Pishchik, V.N., 2022. Plant Development of Early-Maturing Spring Wheat (Triticum aestivum L.) under Inoculation with Bacillus sp. V2026. Plants 11, 1817. 10.3390/plants11141817

68. Mousa, W.K., Ghemrawi, R., Abu-Izneid, T., Al Ramadan, N., Al Sheebani, F., 2024. The design and development of EcoBiomes: Multi-species synthetic microbial consortia inspired by natural desert microbiome to enhance the resilience of climate-sensitive ecosystems. Heliyon 10, e36548. 10.1016/j.heliyon.2024.e36548

69. Müller, L., Müller, D.C., Kammerecker, S., Fluri, M., Neutsch, L., Remus Emsermann, M., Pelludat, C., 2022. Priority Effects in the Apple Flower Determine If the Siderophore Desferrioxamine Is a Virulence Factor for Erwinia amylovora CFBP1430. Applied and Environmental Microbiology 88, e02433–21. 10.1128/aem.02433-21

70. Nawrocki, E.P., Eddy, S.R., 2013. Infernal 1.1: 100-fold faster RNA homology searches. Bioinformatics 29, 2933–2935. 10.1093/bioinformatics/btt509

71. Naylor, D., Coleman-Derr, D., 2018. Drought Stress and Root-Associated Bacterial Communities. Front. Plant Sci. 8. 10.3389/fpls.2017.02223

72. Neilson, J.W., Quade, J., Ortiz, M., Nelson, W.M., Legatzki, A., Tian, F., LaComb, M., Betancourt, J.L., Wing, R.A., Soderlund, C.A., Maier, R.M., 2012. Life at the hyperarid margin: novel bacterial diversity in arid soils of the Atacama Desert, Chile. Extremophiles 16, 553–566. 10.1007/s00792-012-0454-z

73. Pailliè-Jiménez, M.E., Stincone, P., Pereira, J.Q., Santagapita, P.R., Rodrigues, E., Brandelli, A., 2024. Isolation and Characterization of an Antioxidant Aryl Polyene Pigment from Antarctic Bacterium Lysobacter sp. A03. Mol Biotechnol. 10.1007/s12033-024-01132-7

74. Palma, D.E., Gaete, A., López, D., Marcoleta, A.E., Chávez, F.P., Bravo, L.A., Acuña, J.J., Cambiazo, V., Jorquera, M.A., 2025. Microbial Communities in Permafrost, Moraine and Deschampsia antarctica Rhizosphere Soils near Ecology Glacier (King George Island, Maritime Antarctic). Diversity 17, 86. 10.3390/d17020086

75. Parks, D.H., Chuvochina, M., Rinke, C., Mussig, A.J., Chaumeil, P.-A., Hugenholtz, P., 2022. GTDB: an ongoing census of bacterial and archaeal diversity through a phylogenetically consistent, rank normalized and complete genome-based taxonomy. Nucleic Acids Research 50, D785–D794. 10.1093/nar/gkab776

76. Parks, D.H., Imelfort, M., Skennerton, C.T., Hugenholtz, P., Tyson, G.W., 2015. CheckM: assessing the quality of microbial genomes recovered from isolates, single cells, and metagenomes. Genome Res 25, 1043–1055. 10.1101/gr.186072.114

77. Pascale, A., Proietti, S., Pantelides, I.S., Stringlis, I.A., 2020. Modulation of the Root Microbiome by Plant Molecules: The Basis for Targeted Disease Suppression and Plant Growth Promotion. Front. Plant Sci. 10. 10.3389/fpls.2019.01741

78. Pérez Y Terrón, R., González-Montfort, T., Muñoz-Rojas, J., 2014. Antagonismo microbiano asociado a cepas bacterianas provenientes de jitomate (Lycopersicum esculentum Mill) y maíz (Zea Mays). Revista Iberoamericana de Ciencias 1, 53–60.

79. Rauscher, L., Expert, D., Matzanke, B.F., Trautwein, A.X., 2002. Chrysobactin-dependent Iron Acquisition inErwinia chrysanthemi: FUNCTIONAL STUDY OF A HOMOLOG OF THE ESCHERICHIA COLIFERRIC ENTEROBACTIN ESTERASE *. Journal of Biological Chemistry 277, 2385–2395. 10.1074/jbc.M107530200

80. Rodríguez-Pérez, H., Ciuffreda, L., Flores, C., 2021. NanoCLUST: a species-level analysis of 16S rRNA nanopore sequencing data. Bioinformatics 37, 1600–1601. 10.1093/bioinformatics/btaa900

81. Sandy, M., Butler, A., 2011. Chrysobactin Siderophores Produced by Dickeya chrysanthemi EC16. J Nat Prod 74, 1207–1212. 10.1021/np200126z

82. Schöner, T.A., Gassel, S., Osawa, A., Tobias, N.J., Okuno, Y., Sakakibara, Y., Shindo, K., Sandmann, G., Bode, H.B., 2016. Aryl Polyenes, a Highly Abundant Class of Bacterial Natural Products, Are Functionally Related to Antioxidative Carotenoids. ChemBioChem 17, 247–253. 10.1002/cbic.201500474

83. Schwengers, O., Jelonek, L., Dieckmann, M.A., Beyvers, S., Blom, J., Goesmann, A., 2021. Bakta: rapid and standardized annotation of bacterial genomes via alignment-free sequence identification. Microb Genom 7, 000685. 10.1099/mgen.0.000685

84. Shima, S., Matsuoka, H., Iwamoto, T., Sakai, H., 1984. ANTIMICROBIAL ACTION OF ε-POLY-L- LYSINE. J. Antibiot. 37, 1449–1455. 10.7164/antibiotics.37.1449

85. Shouman, H., Said, H.S., Kenawy, H.I., Hassan, R., 2023. Molecular and biological characterization of pyocyanin from clinical and environmental Pseudomonas aeruginosa. Microbial Cell Factories 22, 166. 10.1186/s12934-023-02169-0

86. Solórzano-Acosta, R.A., Quispe, K.R., 2024. Assessing the role of field isolated Pseudomonas and Bacillus as growth-promoting rizobacteria on avocado (Persea americana) seedlings. Journal of Sustainable Agriculture and Environment 3, e12114. 10.1002/sae2.12114

87. Subramanian, P., Kim, K., Krishnamoorthy, R., Mageswari, A., Selvakumar, G., Sa, T., 2016. Cold Stress Tolerance in Psychrotolerant Soil Bacteria and Their Conferred Chilling Resistance in Tomato (Solanum lycopersicum Mill.) under Low Temperatures. PLoS One 11, e0161592. 10.1371/journal.pone.0161592

88. Tian, Y., Liu, Y., Uwaremwe, C., Zhao, X., Yue, L., Zhou, Q., Wang, Y., Tran, L.-S.P., Li, W., Chen, G., Sha, Y., Wang, R., 2023. Characterization of three new plant growth-promoting microbes and effects of the interkingdom interactions on plant growth and disease prevention. Plant Cell Rep 42, 1757–1776. 10.1007/s00299-023-03060-3

89. Vaser, R., Sikic, M., 2021. Time- and memory-efficient genome assembly with Raven. Nature Computational Science 1, 1–5. 10.1038/s43588-021-00073-4

90. Visca, P., Imperi, F., Lamont, I.L., 2007. Pyoverdine siderophores: from biogenesis to biosignificance. Trends in Microbiology 15, 22–30. 10.1016/j.tim.2006.11.004

91. Wang, L., Zhang, C., Zhang, J., Rao, Z., Xu, X., Mao, Z., Chen, X., 2021. Epsilon-poly-L-lysine: Recent Advances in Biomanufacturing and Applications. Front. Bioeng. Biotechnol. 9. 10.3389/fbioe.2021.748976

92. Wang, Y., Liu, H., Shen, Z., Miao, Y., Wang, J., Jiang, X., Shen, Q., Li, R., 2022. Richness and antagonistic effects co-affect plant growth promotion by synthetic microbial consortia. Applied Soil Ecology 170, 104300. 10.1016/j.apsoil.2021.104300

93. Wang, Z., Hu, X., Solanki, M.K., Pang, F., 2023. A Synthetic Microbial Community of Plant Core Microbiome Can Be a Potential Biocontrol Tool. J. Agric. Food Chem. 71, 5030–5041. 10.1021/acs.jafc.2c08017

94. Wang, Z., Piao, Y., Zhang, F., Hu, Y., Zeng, J., Nan, J., 2020. Promoting Effects on Watermelon and Fermentation Optimization of Plantibacter sp. WZW03. J Plant Growth Regul 39, 970–980. 10.1007/s00344-019-10037-8

95. Wani, P.A., Khan, M.S., 2010. *Bacillus* species enhance growth parameters of chickpea (*Cicer arietinum* L.) in chromium stressed soils. Food and Chemical Toxicology 48, 3262–3267. 10.1016/j.fct.2010.08.035

96. Wei, H.-L., Zhang, L.-Q., 2006. Quorum-sensing system influences root colonization and biological control ability in Pseudomonas fluorescens 2P24. Antonie Van Leeuwenhoek 89, 267–280. 10.1007/s10482-005-9028-8

97. Wick, R.R., Holt, K.E., 2019. Benchmarking of long-read assemblers for prokaryote whole genome sequencing. F1000Res 8, 2138. 10.12688/f1000research.21782.4

98. Wick, R.R., Judd, L.M., Cerdeira, L.T., Hawkey, J., Méric, G., Vezina, B., Wyres, K.L., Holt, K.E., 2021. Trycycler: consensus long-read assemblies for bacterial genomes. Genome Biology 22, 266. 10.1186/s13059-021-02483-z

99. Yadav, A.N., Kour, D., Yadav, N., 2023. Microbes as a gift from God. Journal of Applied Biology & Biotechnology. 10.7324/JABB.2023.157095

100. Yadav, A.N., Sachan, S.G., Verma, P., Tyagi, S.P., Kaushik, R., Saxena, A.K., 2015. Culturable diversity and functional annotation of psychrotrophic bacteria from cold desert of Leh Ladakh (India). World J Microbiol Biotechnol 31, 95–108. 10.1007/s11274-014-1768-z

101. Yavarian, S., Jafari, P., Akbari, N., Feizabadi, M.M., 2021. Selective screening and characterization of plant growth promoting bacteria for growth enhancement of tomato, Lycopersicon esculentum. Iran J Microbiol 13, 121–129. 10.18502/ijm.v13i1.5502

102. Ye, R., Xu, Hengyi, Wan, C., Peng, S., Wang, L., Xu, Hong, Aguilar, Z.P., Xiong, Y., Zeng, Z., Wei, H., 2013. Antibacterial activity and mechanism of action of ε-poly-l-lysine. Biochemical and Biophysical Research Communications 439, 148–153. 10.1016/j.bbrc.2013.08.001

103. Zuluaga, M.Y.A., Milani, K.M.L., Miras-Moreno, B., Lucini, L., Valentinuzzi, F., Mimmo, T., Pii, Y., Cesco, S., Rodrigues, E.P., Oliveira, A.L.M. de, 2021. Inoculation with plant growth-promoting bacteria alters the rhizosphere functioning of tomato plants. Applied Soil Ecology 158, 103784. 10.1016/j.apsoil.2020.103784

